# Podocyte lineage marker expression is preserved across Wilms tumor subtypes and enhanced in tumors harboring the SIX1/2 p.Q177R mutation

**DOI:** 10.1101/2022.11.02.514796

**Authors:** Matthew J. Stevenson, Sabrina K. Phanor, Urvi Patel, Stephen S. Gisselbrecht, Martha L. Bulyk, Lori L. O’Brien

## Abstract

Wilms tumors present as an amalgam of varying proportions of three tissues normally located within the developing kidney, one being the multipotent nephron progenitor population. While incomplete differentiation of the nephron progenitors is widely-considered the underlying cause of tumor formation, where this barrier occurs along the differentiation trajectory and how this might promote therapeutic resistance in high-risk blastemal-predominant tumors is unclear. Comprehensive integrated analysis of genomic datasets from normal human fetal kidney and high-risk Wilms tumors has revealed conserved expression of genes indicative of podocyte lineage differentiation in tumors of all subtypes. Comparatively upregulated expression of several of these markers, including the non-canonical WNT ligand *WNT5A*, was identified in tumors with the relapse-associated mutation SIX1/2 p.Q177R. These findings highlight the shared progression of cellular differentiation towards the podocyte lineage within Wilms tumors and enhancement of this differentiation program through promotion of non-canonical WNT/planar cell polarity signaling in association with SIX1/2 p.Q177R.

## INTRODUCTION

Wilms tumor, the most common childhood kidney cancer is typically diagnosed between 2-5 years of age and accounts for an estimated 5% of all cancers in patients under the age of 14 (Steliarova-Foucher et al 2017, Breslow et al 2006, Hol et al 2018). The histological composition of Wilms tumors resembles that of the normal developing kidney, being comprised of varying degrees of blastemal, epithelial, and stromal tissues. Each of these tissues is thought to represent distinct compartments within the developing kidney: 1) nephron progenitor cells (NPCs), the multipotent progenitor population that gives rise to all epithelial cell types of the nephron, 2) differentiating/differentiated tubules, and 3) interstitial cells (ICs), respectively (reviewed in Rivera and Huber, 2005). Beyond the morphological similarities, numerous microarray and transcriptomic analyses have revealed gene expression signatures in Wilms tumors resembling those of normal pre- and post-induction NPCs with varying degrees of differentiation, altogether implying stalled nephrogenesis underlies formation of these tumors (Li et al 2002, Fukuzawa et al 2017, Wegert et al 2015, Walz et al 2015, Gadd et al 2012, Gadd et al 2017, Young et al 2018, Trink et al 2018). Despite the apparent morphological and molecular similarities to normal human fetal kidney (hFK) and the low mutational burden characteristic of Wilms tumors and pediatric malignancies in general compared to that of adult cancers, the genetic drivers and molecular mechanisms underlying the genesis of these tumors remain elusive (Wegert et al 2015, Grobner et al 2018, Kandoth et al 2013).

Current treatment of Wilms tumor is generally considered a success with a five-year relative survival over 90% (SEER cancer stats 1975-2017), yet high-risk blastemal predominant tumors continue to present a challenge to therapeutic intervention. From 2001-2012, 20% of patients with blastemal-predominant tumors treated according to International Society of Paediatric Oncology protocols relapsed within five years of diagnosis and 95% of relapses were distant metastases (Van den Heuvel-Eibrink et al 2015). In patients treated according to Children’s Oncology Group guidelines, 88% of relapses of stage III tumors occurred within the first two years after diagnosis, including five of seven blastemal tumors analyzed in that study (Fernandez et al 2018). Five-year overall survival after relapse of favorable histology Wilms tumor (FHWT) is estimated at 60%-70% (Mullen et al 2018), highlighting the need for improved therapies to limit the risk of relapse for all Wilms tumors, but particularly in the case of high-risk blastemal-predominant tumors.

Mutations in NPC-associated transcription factors *SIX1* and *SIX2* (~7%) are most associated with blastemal-predominant tumors (Gadd et al 2017, Walz et al 2015, Wegert et al 2015). Of the variants detected in *SIX1* across several studies, the overwhelming majority were a glutamine to arginine substitution, p.Q177R (hereafter referred to as Q177R). This mutation is significantly associated with relapse having recently been identified in ~13% of relapsed tumors (Gadd et al 2022). Wegert et al provided the first mechanistic investigation of the SIX1-Q177R mutant protein, demonstrating a unique DNA binding motif for SIX1-Q177R in primary Wilms tumor tissue compared to that of wild type SIX1 in a similar tumor. Increased binding of SIX1-Q177R was observed near the *TGFA* gene with corresponding increases in expression of *TGFA* in SIX1-Q177R mutant tumors compared to wild type SIX1-expressing tumors (Wegert et al 2015). More broadly however, the consequences of this altered DNA binding on the fidelity of downstream transcriptional regulatory networks and its potential role in Wilms tumorigenesis and relapse remain unexplored.

Six1 is required for kidney development in the mouse as knockout results in kidney agenesis. Six1 also acts upstream of canonical NPC markers including Six2 and Pax2, placing it near the top of the NPC gene regulatory network hierarchy (Xu et al 2003, Li et al 2003). Comparative investigations of mouse and human kidney development demonstrated prolonged temporal expression of *SIX1*/SIX1 in NPCs through later stages of development in addition to novel regulatory interactions between SIX1 and SIX2 not observed in the mouse (O’Brien et al 2016). This is suggestive of an expanded regulatory role for SIX1 in human nephrogenesis that may underlie its frequent mutation and association with high-risk Wilms tumors.

Over the last several years, pioneering studies of human kidney development through morphological and transcriptomic analyses have characterized the morphogenesis and molecular signatures intrinsic to the progression from proliferative, multipotent NPCs to the complex multifaceted structure of the nephron. These investigations provided the spatial resolution necessary to more precisely distinguish the transcriptomes of cells in the renal vesicle (RV), the earliest epithelial descendent of the NPCs, from polarized constituents of the S-shaped body (SSB) destined to differentiate to tubules or podocytes (Lindström et al 2018a-d, Lindström et al 2021, Hochane et al 2019, Menon et al 2018, Tran et al 2019). Consequently, these enhanced molecular profiles can be repurposed to refine the molecular identity of cells in Wilms tumors.

The goal of this study is to expand upon previous works and define the direct regulatory role of SIX1-Q177R in Wilms tumor within the context of the normal human nephrogenic niche utilizing genomic data available from large-scale studies of Wilms tumors (Walz et al 2015, Wegert et al 2015), in tandem with SIX1 ChIP-seq data from week 17 hFK (wk17hFK) (O’Brien et al 2016). Furthermore, we sought to classify the Wilms tumor transcriptomes within the framework of the developing hFK by merging single cell RNA-seq datasets from week 14 hFK (wk14hFK) and wk17hFKs and integrating the findings from our genomic analyses of Wilms tumor (Lindström et al 2018d, Lindström et al 2021, Tran et al 2019). In doing so, we identify upregulation of genes associated with promotion of the non-canonical WNT/planar cell polarity (PCP) pathway and/or inhibition of the canonical WNT/β-catenin-mediated pathway in SIX1/2-Q177R-expressing Wilms tumors. Moreover, we provide a mechanistic link to *WNT5A* upregulation attributable to enhanced binding affinity of SIX1-Q177R for putative cis-regulatory elements (CREs) and demonstrate conserved expression of podocyte lineage markers in all Wilms tumors analyzed, a subset of which are specifically upregulated in SIX1/2-Q177R-expressing tumors.

## RESULTS

### Putative SIX1-Q177R target genes are associated with distinct biological processes compared to those of SIX1 in Wilms tumor

Analysis of Wilms tumor SIX1 ChIP-seq peaks containing the previously identified SIX1/SIX1-Q177R DNA binding motifs revealed marked differences in putative regulatory target genes of the two proteins (Figure 1A, Supplemental Figure 1A) (Data kindly provided by Dr. Manfred Gessler, Wegert et al 2015). Comparison with wk17hFK SIX1 ChIP-seq showed modest overlap in peak locations, 41% and 31% of wk17hFK peaks overlapped with SIX1 and SIX1-Q177R tumor peaks, respectively (Figure 1B). Genomic Regions Enrichment of Annotations Tool (GREAT) (McLean et al 2010) was used to identify putative target genes in each tumor dataset. Gene ontology analysis using GREAT revealed enrichment in kidney development-related biological processes in those putative target genes shared between the SIX1 and SIX1-Q177R tumors. Despite modest overlap in physical peak locations between the SIX1 tumor and wk17hFK, SIX1 tumor-only targets are nonetheless enriched for kidney developmental processes. Putative target genes exclusive to SIX1-Q177R, however, are associated with distinct biological processes not explicitly related to kidney development. (Figure 1C).

**FIGURE 1:**
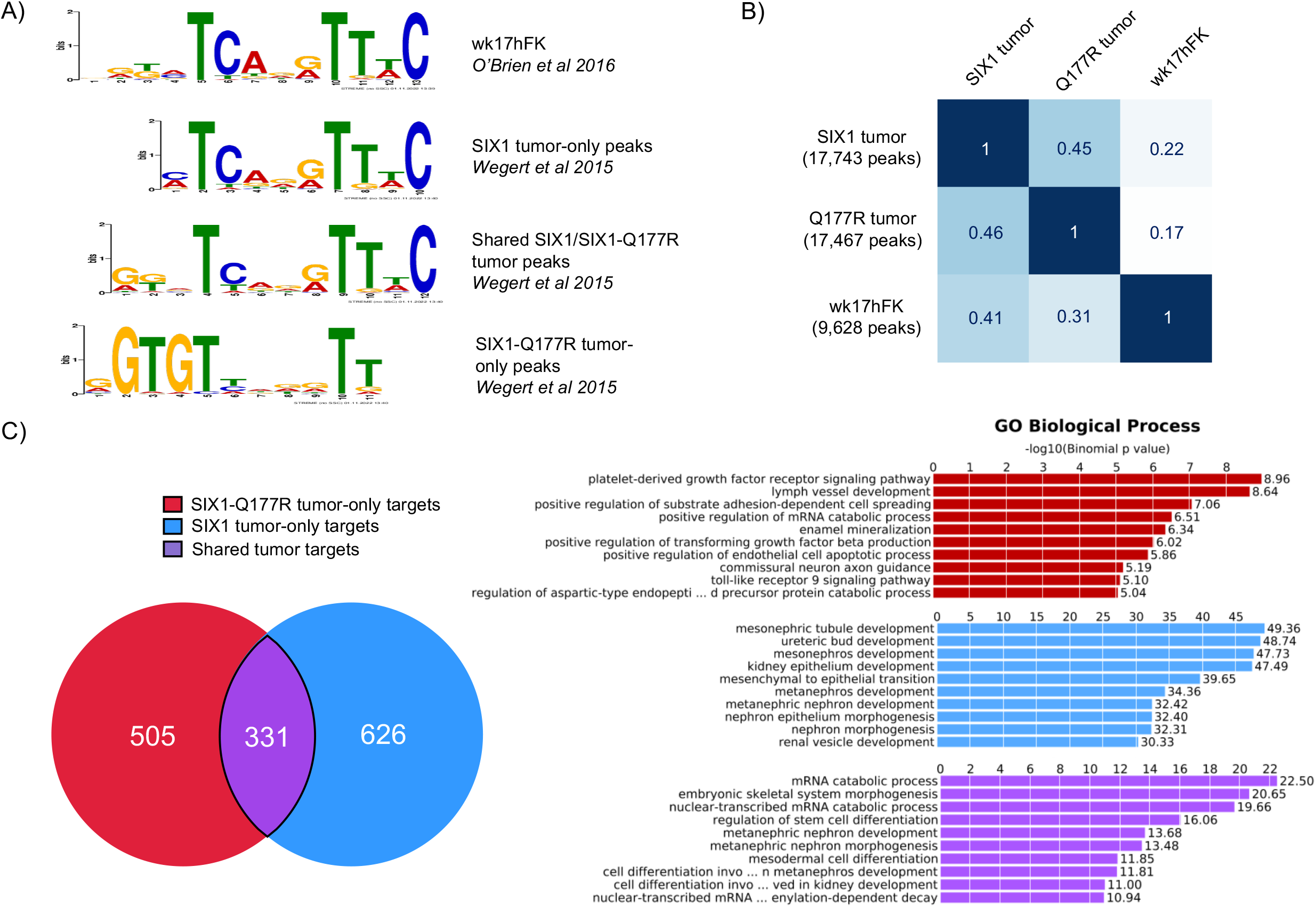
SIX1-Q177R tumor-only ChIP-seq peaks minimally overlap with wk17hFK SIX1 ChIP-seq peaks and putative target genes are associated with biological processes unrelated to kidney development. **A)** SIX1 and SIX1-Q177R DNA binding motifs discovered using the STREME tool (see Experimental Procedures) from Wilms tumor and week 17 human fetal kidney (wk17hFK) SIX1 ChIP-seq datasets (Wegert et al 2015; motifs published previously in that study, O’Brien et al 2016). **B)** Grid displaying the pairwise fraction of SIX1 ChIP-seq peak location overlaps between SIX1 Wilms tumor, SIX1-Q177R Wilms tumor, and wk17hFK. **C)** Venn diagram displaying overlaps of putative target genes identified using motif-enriched peaks in GREAT between SIX1-Q177R Wilms tumor and SIX1 Wilms tumor. Bar charts displaying top 10 most significantly enriched biological processes for shared tumor, SIX1-Q177R tumor-only, and SIX1 tumor-only putative target genes generated using GREAT (FDR < 0.05, significant by both the binomial and hypergeometric tests).

Intriguingly, the MEIS1 DNA binding motif is significantly enriched in all peak sets analyzed (Supplemental Figure 1B). Expression of *Meis1* is restricted to the interstitial lineage in the developing mouse kidney. However, comparative analysis of marker expression by immunofluorescence and *in situ* hybridization in developing hFK revealed overlap of NPC marker *SIX2/SIX2* with *MEIS1/MEIS1* as well as FOXD1/*FOXD1*, another marker used to distinguish interstitial progenitors from NPCs in the mouse (Lindström et al 2018a). Overexpression of either SIX1 or SIX1-Q177R, alongside MEIS1-3xFLAG *in vitro* followed by immunoprecipitation (IP) with a SIX1 antibody resulted in co-IP of MEIS1-3xFLAG (Supplemental Figure 1C). To our knowledge this is the first biochemical evidence of an interaction between SIX1 and MEIS1 in any context. Further investigation is needed to characterize potential regulatory functions of complexes containing these proteins in NPCs.

As the SIX1-Q177R ChIP-seq data was obtained from a single Wilms tumor (Wegert et al 2015), we sought to interrogate the DNA binding preference of the mutant protein in the absence of potentially confounding variables including tissue quality and chemotherapy-induced artifacts. To determine the effect on sequence specificity of the variant, we expressed the reference allele and the Q177R mutant SIX1 homeodomains *in vitro* and assayed their specificities in parallel by protein binding microarrays (PBMs) (Berger et al 2006). The primary and secondary motifs (Badis et al 2009) recognized by both alleles are shown in Figure 2A; a replicate experiment on an independent array yielded qualitatively similar logos. To quantify the binding of each allele to each motif, we calculated “pattern E-scores” (see Methods) for each replicate of each protein to the four patterns shown in Figure 2B. The Q177R allele showed strikingly reduced binding to the reference primary motif and essentially no binding to the reference secondary motif, while the alternate motifs bound by the Q177R allele showed similarly poor binding by the reference allele. These findings validate those of Wegert et al from primary Wilms tumor tissue and confirm the distinct binding specificity of the mutant protein is a direct result of the Q177R mutation.

**FIGURE 2:**
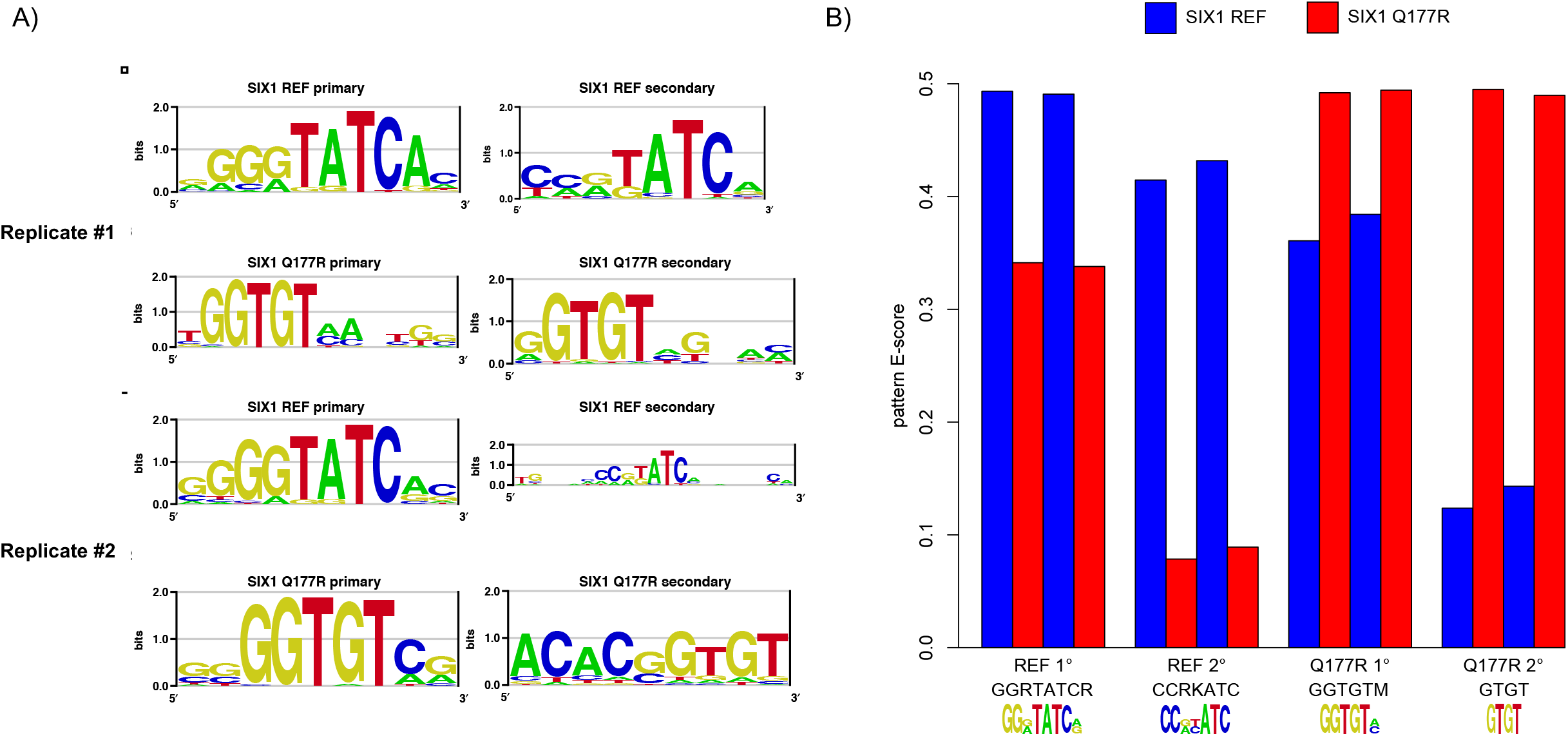
The Q177R variant shifts the sequence preference of SIX1. **A)** Representative logos of the primary binding preferences of the SIX1 reference allele (top) and Q177R variant (bottom), and of the additional (“secondary”) binding specificity not well explained by the primary logos. **B)** Pattern E-scores (see Experimental Procedures) for binding of the SIX1 reference allele (blue) and Q177R variant (red) to each indicated pattern.

### *WNT5A* and other WNT signaling effectors are upregulated in SIX1/2-Q177R Wilms tumors

To further refine the general peak/gene associations, we utilized RNA-seq data from 114 chemotherapy-naïve high-risk FHWTs and diffuse anaplastic Wilms tumors (DAWT) of varying histological classifications that subsequently relapsed, generated as part of the Therapeutically Applicable Research to Generate Effective Treatments (TARGET) program, to pinpoint differentially expressed genes unique to SIX1/2-Q177R tumors. The same Q177R mutation has been identified in SIX2 in Wilms tumors but occurs almost half as frequently as SIX1-Q177R (Wegert et al 2015, Walz et al 2015, Gadd et al 2017). SIX2 is a closely related Six family transcription factor to SIX1, sharing nearly 100% amino acid residue conservation within the DNA binding homeodomain and binds to most of the same genomic sites as SIX1 in hFK (O’Brien et al 2016). Accounting for these similarities between SIX1/SIX2 and to increase the statistical power of our analyses, we grouped SIX1-Q177R and SIX2-Q177R tumors.

Nearly 30% of SIX1/2-Q177R tumors also harbor inactivating mutations in the miRNA processing genes *DROSHA* or *DGCR8* (Wegert et al 2015, Walz et al 2015, Gadd et al 2017). Strikingly, co-occurrence of SIX1/2-Q177R and DROSHA/DGCR8 mutations significantly increased the rates of relapse and death in a synergistic manner compared to tumors with other mutations and tumors with SIX1/2-Q177R or DROSHA/DGCR8 mutations alone (Walz et al 2015). Notably, Wegert et al observed no strong effects on gene expression in DROSHA/DGCR8 mutant tumors, only a significant reduction in miRNA levels (Wegert et al 2015). Due to the scarcity of SIX1/2-Q177R tumors without mutations in miRNA processing genes in this dataset and to best isolate the transcriptional effects attributable to SIX1/2-Q177R, tumors harboring SIX1/2-Q177R with or without miRNA mutations were included in differential expression analysis (SIX1/2miRNA, n=9). Specific histology classification of other tumors used in this analysis was based on prior classification if available (Walz et al 2015, TARGET data matrix https://portal.gdc.cancer.gov/projects), otherwise tumors were included in the group containing mixed histology tumors, generating three additional tumor groups: wild type SIX1-expressing blastemal tumors (Blastemal, n=22), tumors of mixed/epithelial/stromal histology (MIXED/ES, n=56), and DAWT (n=27) (Supplemental Table 1). Of note, although SIX1/2-Q177R is most associated with blastemal histology (Wegert et al 2015, Walz et al 2015), three of the SIX1/2miRNA tumors used in this analysis were classified as mixed histology and one tumor was classified as DAWT (Walz et al 2015, Gadd et al 2017).

Limma-voom was utilized for differential gene expression analysis (Supplemental Figure 2) (Smyth 2005, Law et al 2014, Liu et al 2015). Due to the overwhelming evidence supporting the nephrogenic origin of Wilms tumors (reviewed in Li et al 2021 and Hohenstein et al 2015, Coorens et al 2019), we reasoned an approach focusing on transcripts expressed in all tumor samples could elucidate important deviations along the nephrogenic trajectory between tumors, while also highlighting critical shared intrinsic characteristics of these tumors. Therefore, weakly-expressed genes were stringently filtered to exclude potential false positives due to heterogeneity in tumor microenvironments or sample-to-sample processing variability (genes with counts per million (CPM) < 2 in any sample were removed). Unsupervised hierarchical clustering by Pearson’s correlation coefficient using normalized and scaled gene expression values illustrated a high degree of transcriptional similarity between the majority of tumors in the dataset, regardless of histological classification (Figure 3A). Moreover, tumors with Chromosome 1 q21-q23 gain, recently identified in 75% of tumors after relapse (Gadd et al 2022), did not noticeably cluster together suggesting the magnitude of transcriptional changes associated with this copy number gain are minimal or heterogeneous and will not compromise the interpretations of our differential gene expression analyses. As has been noted previously, most SIX1/2miRNA tumors clustered together (Wegert et al 2015, Walz et al 2015).

**FIGURE 3:**
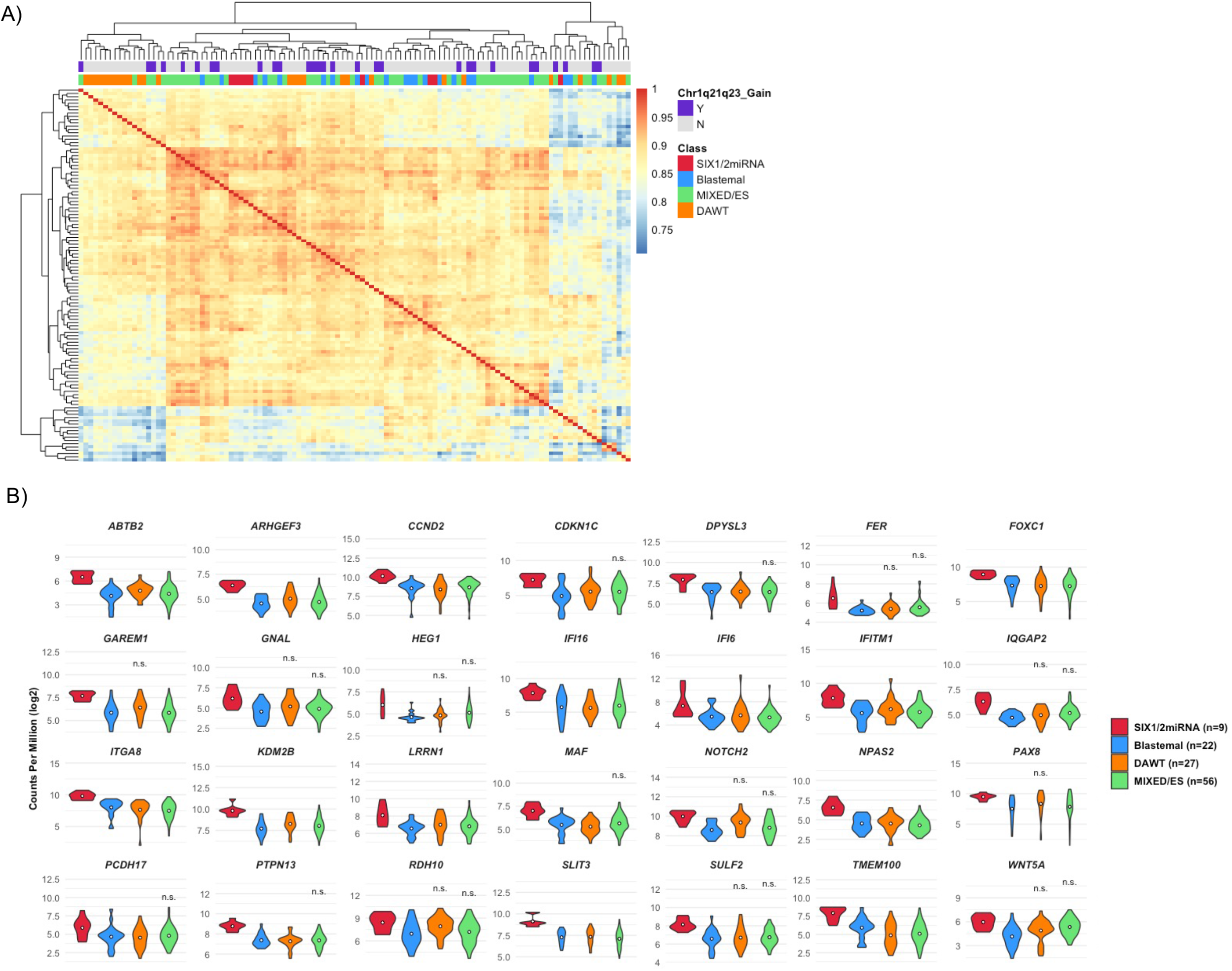
Several genes upregulated in SIX1/2miRNA tumors have functions related to enhancement of non-canonical WNT/PCP signaling and/or inhibition of canonical WNT/β-catenin-mediated signaling. **A)** Heatmap displaying results of unsupervised hierarchical clustering of all tumors used in RNA-seq differential gene expression analysis by Pearson’s correlation coefficient (see Experimental Procedures). Horizontal bar directly above heatmap is color-coded to identify differential gene expression group (class) of each tumor as indicated in legend. Uppermost horizontal bar above heatmap is color-coded to indicate presence (purple) or absence (gray) of Chromosome 1 q21-q23 amplification status of each tumor. **B)** Violin plots showing the distributions of log_2_ counts per million (CPM) of genes significantly upregulated in SIX1/2miRNA tumors compared to Blastemal tumors. Dot within each group plot represents the mean. Unless indicated by n.s. (not significant), the log_2_ fold change of the SIX1/2miRNA group was > 1.5 and adjusted p-value < 0.05 compared to that tumor group.

Few genes were identified as differentially expressed between Blastemal and MIXED/ES groups using a less stringent fold change cutoff, eight genes upregulated and 20 genes downregulated (log_2_ fold change > |1|, adj. p < 0.05) (Supplemental Figure 3A, Supplemental File 1). Compared to Blastemal tumors, 28 genes were significantly upregulated in SIX1/2miRNA tumors, 14 of which were also significantly upregulated compared to MIXED/ES tumors (log_2_ fold change > 1.5, adj. p <0.05) (Figure 3B, Supplemental Figure 3A, Supplemental File 1). Remarkably, the expression levels of well-characterized NPC markers including *SIX1, SIX2, SALL1, MEOX1, PAX2, LYPD1*, and *CRABP2* were not significantly different between the four groups (Supplemental File 1). Thus, blastemal-predominant tumors do not appear to represent an augmentation of the pre-induction NPC transcriptional regulatory identity compared to tumors of other histology. *CCND2*, encoding cyclin D2, was significantly upregulated in SIX1/2miRNA tumors compared to both Blastemal and DAWT groups and was upregulated 1.46 log_2_ fold compared to MIXED/ES tumors, similar to findings of other studies (Wegert et al 2015, Walz et al 2015). Nine genes were significantly upregulated in SIX1/2miRNA tumors compared to all groups, including known NPC markers *TMEM100* and *ITGA8*. This suggests the sustained proliferation reflected by enhanced *CCND2* expression in SIX1/2miRNA tumors may be governed by non-transcriptional mechanisms including cell adhesion. Several genes were significantly downregulated in SIX1/2miRNA tumors as well and will be addressed in a later section, however we chose to focus on upregulated genes for initial follow-up analyses (Supplemental Figure 3B, Supplemental File 1).

Functional similarities among the upregulated SIX1/2miRNA genes were discovered through literature searches, revealing an enrichment in protein functions either promoting non-canonical WNT/PCP signaling and/or antagonizing canonical WNT/β-catenin signaling. Positive regulators of non-canonical WNT/PCP signaling include non-canonical WNT ligand WNT5A (Qian et al 2007, Nishita et al 2010), IQGAP2 through interaction with Cdc42/Rac1 (Logue et al 2011, Ozdemir et al 2018, Fukata et al 2002), ARHGEF3 through activation of RhoA (D’Amato et al 2015, You et al 2021), and FOXC1 through direct regulation of *WNT5A* expression (Han et al 2018). Negative regulators of canonical WNT/β-catenin signaling include WNT5A through interaction with the receptor Ror2 (Mikels and Nusse 2006), KDM2B through independent demethylation of β-catenin and transcriptional repression of β-catenin target genes (Lu et al 2015, Ladinovich et al 2020), as well as SLIT3 and IQGAP2 as the expression of both has been associated with decreased β-catenin target gene activation or decreased nuclear β-catenin, respectively (Kim et al 2018, Ng et al 2018, Deng et al 2016). These findings suggest imbalance in WNT/β-catenin signaling in favor of non-canonical WNT/PCP may contribute to the enhanced aggressive nature of SIX1/2miRNA tumors. Of particular interest as it relates to the increased rates of relapse of SIX1/2-Q177R tumors is *WNT5A*, as it has been associated with increased resistance to chemotherapeutics in ovarian and breast cancer cells (Peng et al 2011, Hung et al 2014).

### SIX1 and SIX1-Q177R enhance transcription from proximal and distal *WNT5A* CREs *in vitro* and SIX1-Q177R binds the *WNT5A* proximal CRE with higher affinity than SIX1

Again, leveraging the available SIX1 ChIP-seq data, we identified peaks shared between SIX1 and SIX1-Q177R in Wilms tumors at proximal and distal regions near the *WNT5A* locus. High DNA sequence conservation suggests these sites may represent CREs. Moreover, these peaks did not appear in the wk17hFK dataset (Figure 4A). Apart from *WNT5A*, proximal and/or distal SIX1-Q177R or shared tumor peaks were identified within candidate CREs as predicted by ENCODE or GeneHancer for other upregulated SIX1/2miRNA genes including *KDM2B, CDKN1C, TMEM100, ARHGEF3*, and *FOXC1* (Supplemental Figure 4A) (Abascal et al 2020, Fishilevich et al 2017).

**FIGURE 4:**
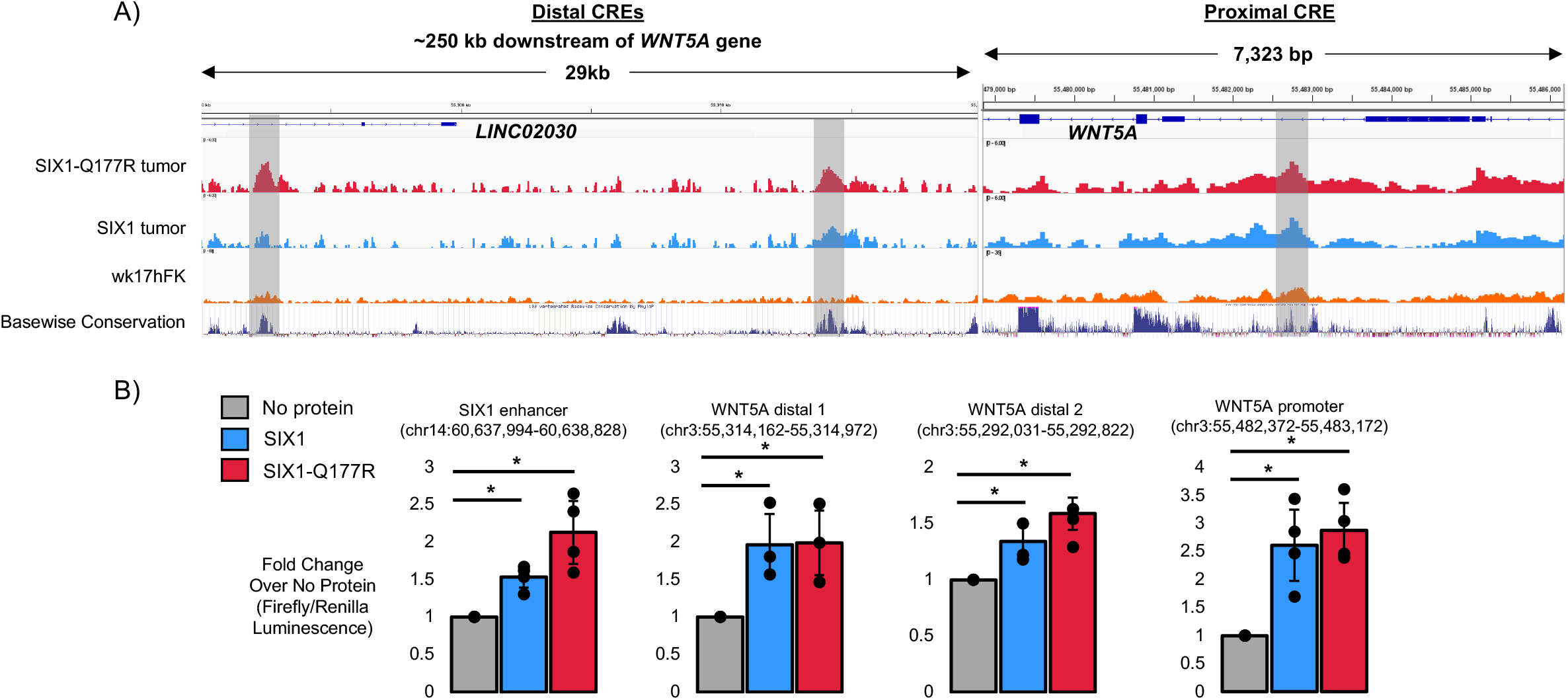
SIX1 and SIX1-Q177R bound highly-conserved putative proximal and distal cis-regulatory elements (CREs) for *WNT5A* in Wilms tumors and both proteins enhanced expression from each CRE in *in vitro* luciferase enhancer assays. **A)** IGV genome browser snapshots of genomic regions containing putative proximal and distal *WNT5A* CREs displaying SIX1-Q177R tumor, SIX1 tumor, and wk17hFK SIX1 ChIP-seq tracks along with UCSC genome browser basewise conservation track. Shaded regions indicate positions of shared SIX1 and SIX1-Q177R ChIP-seq peaks in Wilms tumors. **B)** Bar graphs displaying results from the luciferase enhancer assays for each of the indicated DNA elements tested using “No protein” condition as control, * = p < 0.05, n ≥ 3 biological replicates per condition.

To explore the potential regulation of gene expression by SIX1-Q177R through the *WNT5A* CREs, luciferase activity was measured from minimal promoter constructs containing each putative regulatory element in MCF-7 cells while overexpressing either SIX1 or SIX1-Q177R alongside the cofactor EYA1. As shown in Figure 4B, both wild type and mutant SIX1 drove significant expression of luciferase from the proximal DNA element and both distal DNA elements. These data support the regulation of *WNT5A* expression by both wild type SIX1 and SIX1-Q177R. Nevertheless, overexpression of the proteins in these assays could mask subtle, yet biologically significant differences in their regulatory activities when expressed at normal physiological levels.

The SIX1-Q177R mutation is almost exclusively heterozygous in Wilms tumors and in the heterozygous context, both wild type *SIX1* and *SIX1-Q177R* alleles were found to be expressed at similar levels (Walz et al 2015, Wegert et al 2015). Accordingly, differences in binding affinity at peaks shared between the two proteins could account for aberrant gene expression *in vivo*. To interrogate this possibility, we carried out electrophoretic mobility shift assays (EMSAs) using purified wild type SIX1 or SIX1-Q177R protein expressed in *E. coli* and biotinylated oligonucleotide probes containing the putative DNA binding sequence found within the *WNT5A* proximal peak which is highly congruent to the primary SIX1-Q177R motif discovered in the ChIP-seq and PBMs (Wegert et al 2015). Shown in Figure 5, SIX1-Q177R binds this sequence with an estimated 10-fold higher affinity compared to SIX1. Furthermore, changing the guanine to an adenine at the nucleotide position that appears to be preferred by SIX1-Q177R (Wegert et al 2015, Figure 1A, Figure 2) resulted in a loss of affinity of SIX1-Q177R with a concomitant, albeit modest increase in affinity of SIX1. Therefore, under conditions of equal expression of SIX1 and SIX1-Q177R, SIX1-Q177R likely outcompetes its wild type counterpart for binding to this CRE, promoting aberrant expression of *WNT5A*. That SIX1-Q177R binds the “mutated” probe in which the DNA sequence aligns with the wild type SIX1 motifs from the ChIPseq and PBMs with similar affinity to SIX1 is somewhat contradictory to the PBM results shown in Figure 2B (O’Brien et al 2016, Wegert et al 2015). However, while the PBMs utilized homeodomain protein fragments, full-length proteins were used in the EMSAs. Therefore, intramolecular interactions within the tertiary structure of the full-length peptides might contribute to binding of less-preferred DNA motifs.

**FIGURE 5:**
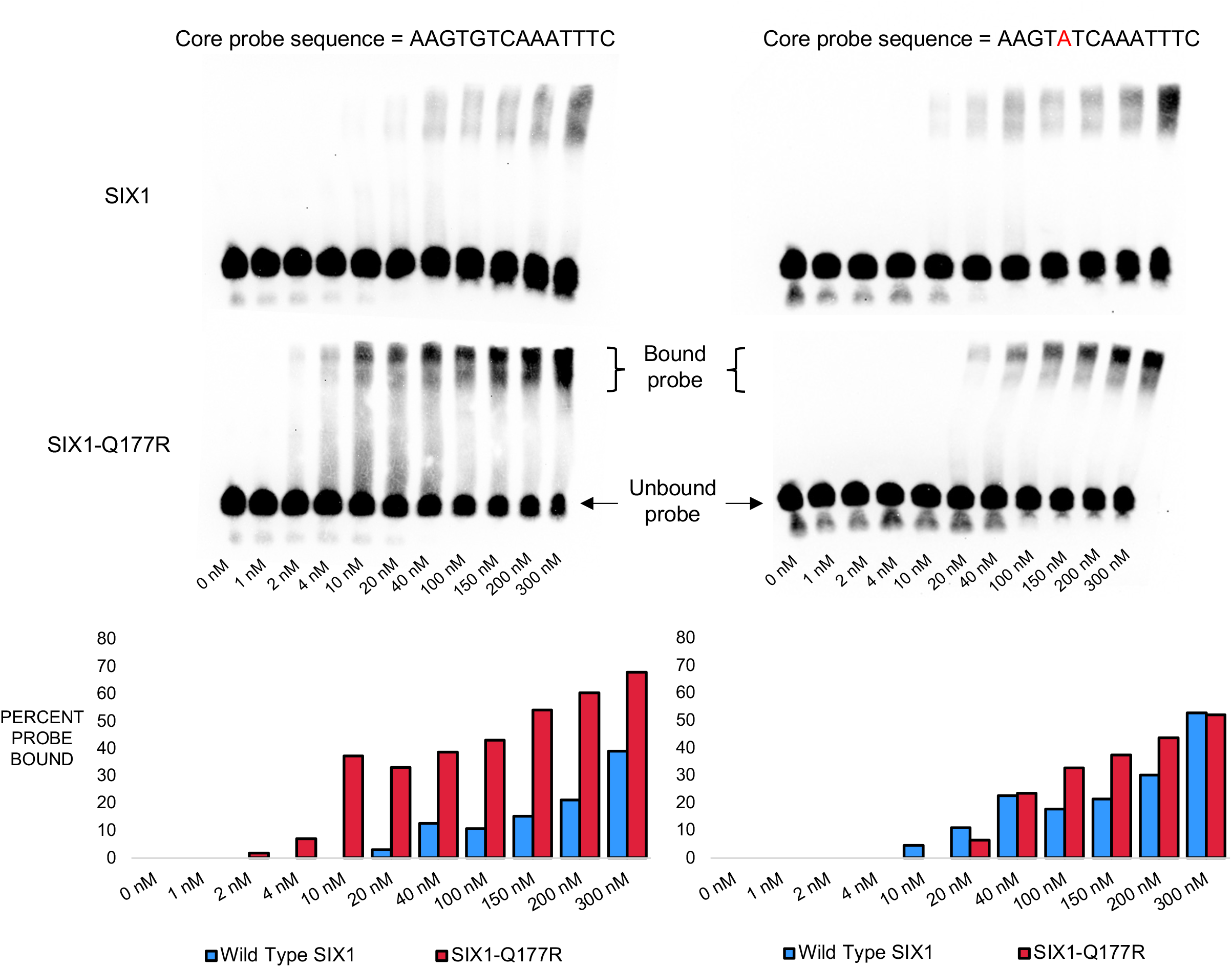
SIX1-Q177R binds core DNA motif sequence from *WNT5A* proximal CRE with higher affinity than SIX1. Top, chemiluminescence images of blots from Electrophoretic Mobility Shift Assays (EMSA) using varying concentrations of purified SIX1 or SIX1-Q177R protein and biotin-labeled oligonucleotide probes with core DNA motif sequence indicated above (see Experimental Procedures). Bottom, bar graphs displaying quantification of “percent probe bound” derived from signal intensities of bound and unbound probe bands (see Experimental Procedures).

### Several upregulated genes in SIX1/2miRNA tumors are characteristic of the podocyte lineage in the developing human kidney

To relate our findings thus far to the normal human nephrogenic niche, wk14hFK and wk17hFK single cell RNA-seq datasets (GSE112570, GSE139280, GSE124472 (only sample GSM3534656)) were integrated using Seurat to generate a powerful reference with which to assess the expression and localization of the differentially expressed Wilms tumor genes (Supplemental Figure 5A) (Hao and Hao et al 2021, Stuart and Butler et al 2019, Butler et al 2018, Satija and Farrell et al 2015). These datasets were derived from specific cortical isolation procedures of hFK, ensuring enrichment of predominantly cortical nephrogenic tissue (Lindström et al 2018d, Lindström et al 2021, Tran et al 2019). Consisting of more than 30,000 cells, UMAP dimensionality generated 20 distinct clusters. Using the same sets of markers for cluster annotation as used in the data source publications, consistent clusters were identified in the integrated dataset (Supplemental Figure 5B, Supplemental File 2). In addition, unsupervised hierarchical clustering by Pearson’s correlation coefficient using average z-scores for each gene across clusters revealed cluster similarities which, in combination with the known cluster marker genes, was used to guide the merging of individual clusters to generate the broad cluster identities shown in Figure 6A (Supplemental Figure 5C).

**FIGURE 6:**
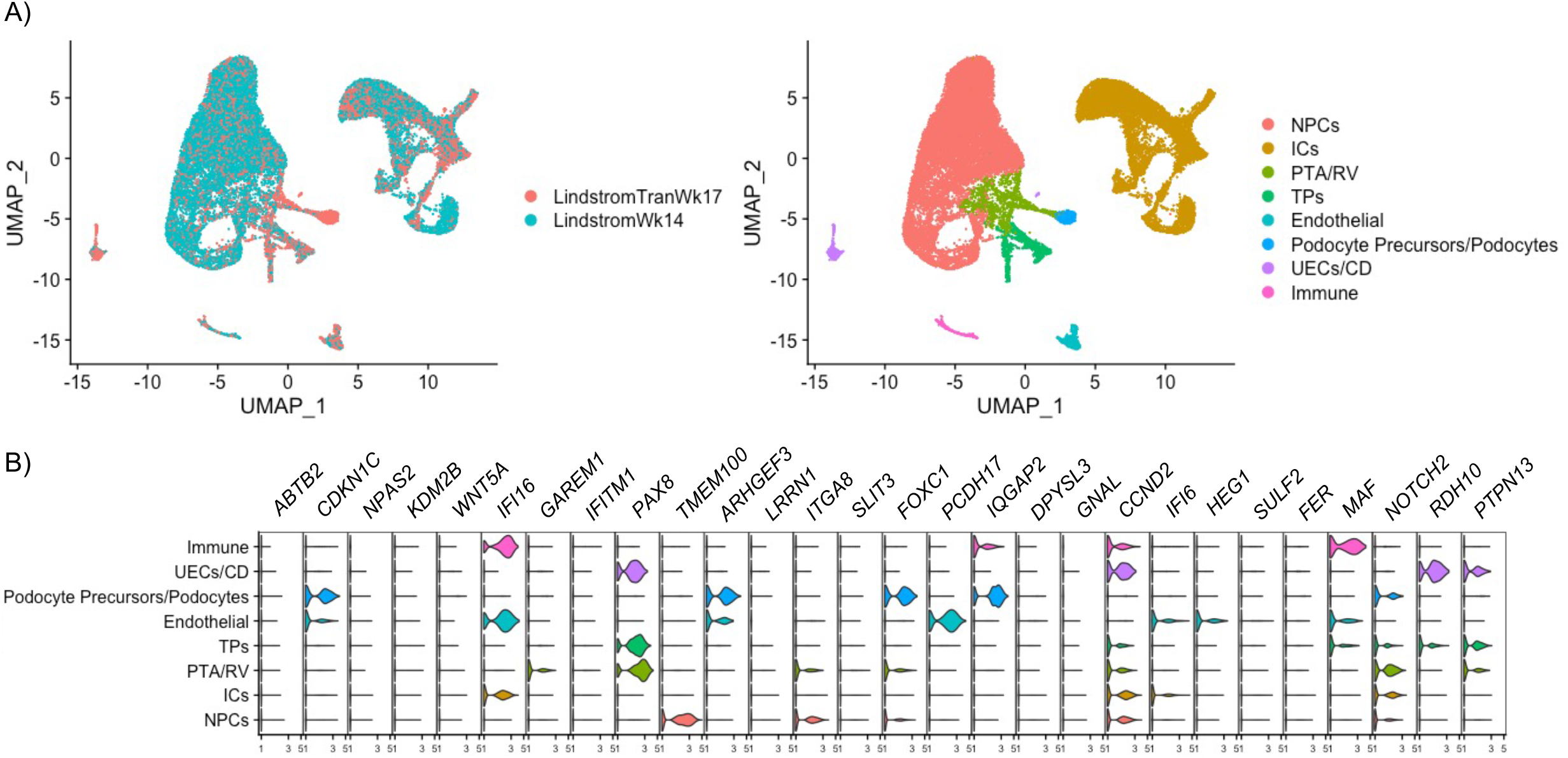
Several genes significantly upregulated in SIX1/2miRNA Wilms tumors show podocyte-enriched expression pattern in developing human kidney. **A)** Left, UMAP dimensional reduction plot showing clustering of cells from LindstromTranWk17 (GSE112570 and GSE124472 (only sample GSM3534656)) and LindstromWk14 (GSE139280) human fetal kidney (hFK) single cell RNA-seq datasets after integrated analysis using Seurat (see Experimental Procedures). Right, UMAP dimensional reduction plot showing cell type classification of cell clusters. NPCs = nephron progenitor cells, ICs = interstitial cells, PTA/RV = pretubular aggregate/renal vesicle, UECs/CD = ureteric epithelial cells/collecting duct. **B)** Violin plots displaying the distribution of normalized expression levels of the significantly upregulated genes in SIX1/2miRNA tumors compared to Blastemal tumors (from Figure 3B) in each hFK cell cluster as shown in panel A.

Probing this dataset for the SIX1/2miRNA upregulated genes shown in Figure 3B revealed abundant and podocyte precursor-enriched expression of *CDKN1C, ARHGEF3, FOXC1*, and *IQGAP2* (Figure 6B). Expression of *NOTCH2* also appears in podocyte precursors as well as PTA/RV alongside *PAX8*, a canonical marker of NPC differentiation. Notably, *WNT5A* is also highly expressed in a small population of podocyte precursors, indicating *WNT5A* expression is associated with podocyte differentiation (Supplemental Figure 6). Although *KDM2B* does not appear strongly in this dataset, *in situ* hybridization in hFK has illustrated high expression in RV/SSB structures (Lindström et al 2018d). Interrogation of the genes significantly downregulated in SIX1/2miRNA tumors compared to both Blastemal and MIXED/ES tumors reveal enrichment of genes with IC-specific expression patterns, likely indicative of smaller proportions of stroma/ICs in the SIX1/2miRNA tumors (Supplemental Figure 6). The expression pattern of the upregulated genes in differentiating structures of the developing human kidney is indicative of a farther progression along the podocyte differentiation trajectory in SIX1/2miRNA tumors than has been characterized previously (Walz et al 2015, Wegert et al 2015).

### Numerous genes exhibiting podocyte-enriched expression pattern in the developing kidney are expressed in all Wilms tumors analyzed

Due to our stringent filtering of weakly-expressed genes described earlier, we can infer with high confidence that the genes passing this filtering step are expressed at appreciable levels in all tumor samples. As such, we examined these genes more closely, regardless of significant differential expression to gain additional insight into the correlation of the Wilms tumor transcriptome with that of the developing hFK. Removal of human housekeeping genes (Eisenberg and Levanon 2013) and intersection of the remaining genes with markers of all clusters in the integrated single cell RNA-seq dataset generated a list of 1,149 genes (Supplemental File 2). Analysis of average z-scores for each gene across all cell clusters shows the largest proportions of genes are expressed predominantly in endothelial cells, immune cells, and podocyte precursors in the hFK (Figure 7A). Gene Ontology analysis using the Database for Annotation, Visualization, and Integrated Discovery (DAVID) revealed significant enrichment of biological processes related to angiogenesis, cell migration, and cell adhesion (Figure 7B) (Sherman et al 2022, Huang et al 2009). That large fractions of these genes are associated with endothelial and immune cells is unsurprising, as there is evidence for the presence of both endothelium and tumor-associated macrophages in Wilms tumors (Skoldenberg et al 2001, Ghanem et al 2003, Liou 2013, Vakkila 2016, Tian 2020). The prevalence of podocyte-precursor specific genes is consistent with the observation within the SIX1/2miRNA upregulated gene set. To characterize the podocyte lineage relationship further, we examined the expression of genes identified by Tran et al as signatures of cells at early (EP) and late (LP) stages along the podocyte differentiation trajectory. The hFK expression patterns of those genes that were expressed in all Wilms tumors analyzed and were also identified as EP and LP genes by Tran et al are shown in Figure 7C (Tran et al 2019). The expression of selected genes from that list within each tumor group are shown in Figure 7D, including canonical podocyte markers *MAFB, PODXL*, and *SYNPO*. These data illustrate that high-risk Wilms tumors of all histological subtypes recapitulate the continuum of podocyte specification at the transcriptional level.

**FIGURE 7:**
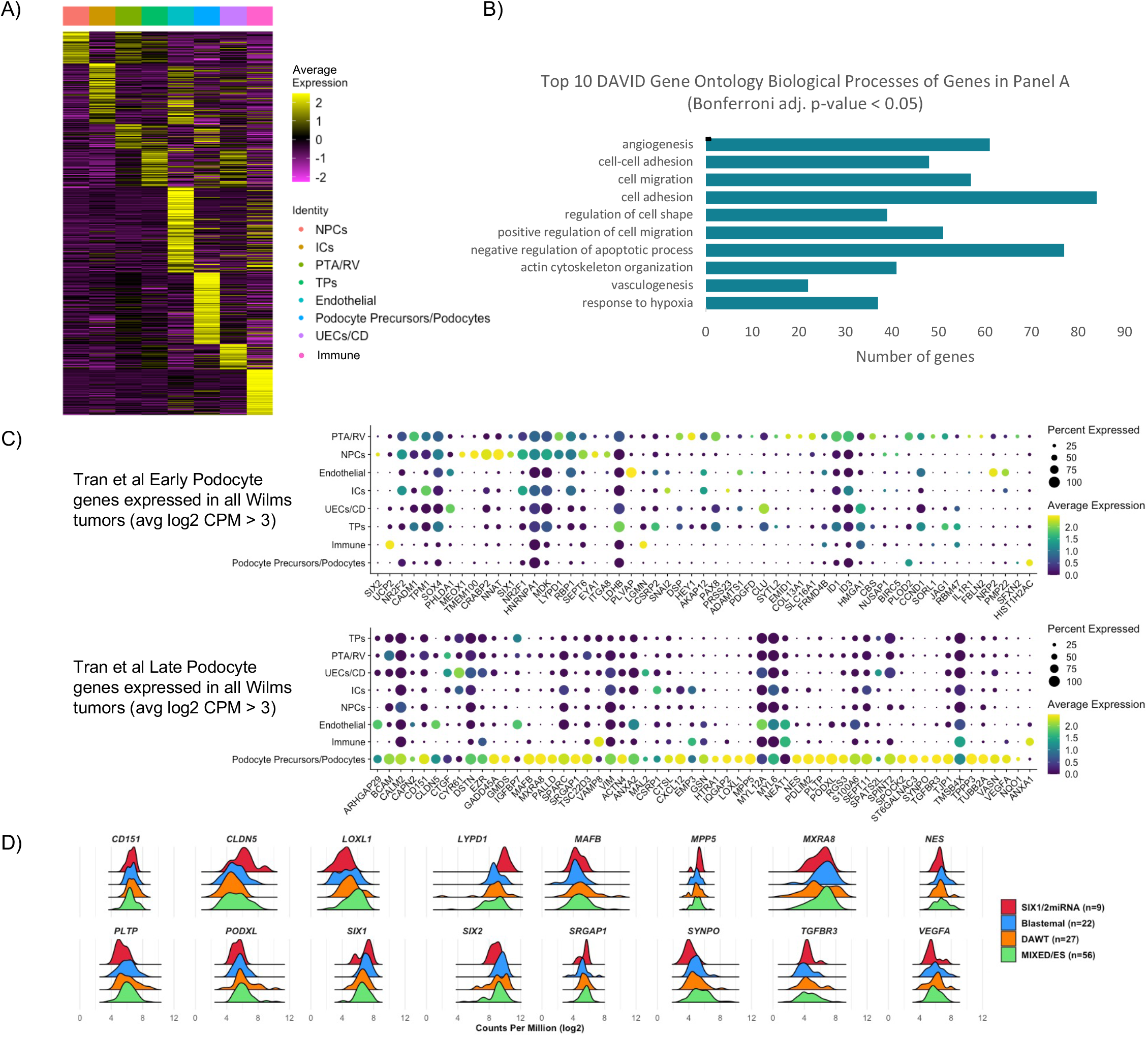
Numerous genes expressed in all Wilms tumor groups show podocyte-enriched expression pattern in developing human kidney, recapitulating both early and late signatures of podocyte lineage differentiation. **A)** Heatmap displaying z-scores of average expression values of all cells within each human fetal kidney (hFK) cell cluster for the 1,149 genes expressed in all Wilms tumor groups that were also identified as markers of any of the hFK cell clusters (see Experimental Procedures). **B)** Bar chart showing the number of genes from Panel A annotated in each of the shown Gene Ontology Biological Process terms identified using the Database for Annotation, Visualization and Integrated Discovery (DAVID). **C)** Dot plots showing scaled average expression values of the indicated genes in all cells within the indicated hFK cell clusters and the percentage of cells within each cluster expressing each gene. Genes displayed were derived from Tran et al early podocyte (top) and late podocyte (bottom) gene signatures (Tran et al, 2019). **D)** Density ridgeline plots showing the distribution of log_2_ CPM expression values of the indicated genes derived from Panel C in all Wilms tumor subgroups.

## DISCUSSION

Here we have integrated genomic datasets from pre-chemotherapy high-risk Wilms tumors and developing hFK to demonstrate the commonality of a podocyte-like gene expression signature in Wilms tumors of all histological subtypes and the augmentation of podocyte-specific gene expression in SIX1/2miRNA tumors. Wilms tumors have been characterized generally as arising from aberrant kidney development, yet until the availability of single cell RNA sequencing the field has lacked the necessary resolution to place Wilms tumors along the normal human nephrogenic differentiation trajectory. Our findings appear to contradict those of prior studies in which the authors observed upregulation of genes associated with pre-induction metanephric mesenchyme in SIX1/2-Q177R tumors (Wegert et al 2015, Walz et al 2015), as well as a broader enrichment of pre-induction NPC gene expression with accompanying low post-induction gene expression in most FHWTs (Gadd et al 2017). However, these works preceded the use of single cell RNA sequencing in the context of human fetal kidney development beginning in 2018.

Subsequently, genes once characterized almost exclusively as pre-induction NPC markers are now known to be expressed in differentiating structures including the PTA and epithelialized RV. This includes *SIX2, CITED1, TMEM100*, and *MEOX1*, and even *SIX1* in the proximal SSB localized to the podocyte precursors (Lindström et al 2018a, Lindström et al 2018c). This raises the likelihood that SIX1 plays a meaningful functional role in the gene regulatory networks governing podocyte lineage specification. Thus, the persistent expression of these genes in Wilms tumors does not necessarily reflect a proliferative, naïve NPC-like state. In fact, a study using serial transplantation of Wilms tumor xenografts suggested cells resembling induced epithelium were the primary culprit of propagation in these tumors (Shukrun et al 2014).

Podocytes, highly specialized epithelial derivatives of NPCs, form extensive mesh-like networks with one another in which cellular protrusions termed “foot processes” interdigitate and surround the glomerular capillaries to filter the incoming blood. While it is well-understood that podocytes derive from the NPCs, the temporal relationship and transcriptional similarity between these cell populations was only recently uncovered. Lindström et al used time-lapse imaging of the developing mouse kidney to demonstrate that the last committed NPCs integrating into the RV contribute to the podocyte lineage. Pseudotime temporal analysis of single-cell RNA-seq data from nephrogenic hFK tissue reinforced those findings, indicating podocytes are the most closely related descendent of NPCs and arise from a distinct differentiation timeline to that of the tubule precursors, so much so that the authors concluded podocytes differentiate directly form committed NPCs (Lindström et al 2018d). Tran et al characterized this intimate relationship in detail using single-cell RNA-seq to distinguish mature podocytes located in the inner cortex of the developing kidney to the NPCs and early podocyte precursors located in the outer cortex of the kidney. This analysis generated a panel of marker genes distinguishing the EP lineage from the LP lineage. In agreement with Lindström et al’s evidence, approximately half of the EP genes are expressed in NPCs including *SIX1, SIX2, CRABP2, MEOX1, TMEM100*, and *ITGA8* (Tran et al 2019).

Our analyses suggest this podocyte lineage trajectory is broadly conserved in high-risk Wilms tumors, with certain cytoskeletal and cell adhesion attributes potentially enhanced by the SIX1/2-Q177R mutation. Imbalance in WNT pathway signaling in favor of non-canonical WNT/PCP in the context of SIX1/2-Q177R would also favor podocyte lineage progression, as canonical WNT/β-catenin signaling has been demonstrated to inhibit podocyte differentiation in mouse and chick model systems, as well as in human induced pluripotent stem cell (hiPSC)-derived podocytes (Lindström et al 2015, Grinstein et al 2013, Yoshimura et al 2019). Supported by the findings of Lindström et al and Tran et al, acquisition of podocyte-like characteristics represents the differentiation path of least resistance. Moreover, the specialized traits of podocytes would likely be advantageous for tumor survival and propagation. Podocytes require a complex cytoskeletal architecture for foot-process formation, tight adherence with the underlying matrix, and maintenance of specialized cell junctions, or slit diaphragms, to ensure proper glomerular filtration (reviewed in Welsh and Saleem 2011). While we do not anticipate that the complex architecture of mature podocytes is recapitulated in these tumors, the acquisition of immature podocyte traits coupled with stalled differentiation would support both continued proliferation and cellular advantages within the normal kidney milieu (reviewed in Welsh and Saleem 2011). Developing podocytes also produce the angiogenic growth factor VEGF-A, as do Wilms tumors (Figure 7D), which is thought to attract endothelial progenitor cells to promote vascularization of the developing glomerulus and could feasibly enhance vascularization of Wilms tumors (Kim et al 2019, Eremina et al 2003, Kitamoto and Tomita 1997).

Altogether, the evidence presented here supports a more podocyte-restricted developmental path of SIX1/2-Q177R-expressing cells in Wilms tumor, the characteristics of which may aid in resistance to chemotherapeutic intervention. However, whether this mutation is sufficient for oncogenic transformation in isolation remains unanswered and unlikely. The most common genomic anomaly in Wilms tumor is the loss of imprinting (LOI)/loss of heterozygosity (LOH) at chromosome 11p15 resulting in overexpression of *IGF2*, occurring in 75-80% of tumors (Gadd et al 2017, Wegert et al 2015). *IGF2* was the most highly expressed gene in our differential expression analysis (log_2_ CPM = ~15). Therefore, synergism between SIX1/2-Q177R and *IGF2* overexpression or miRNA processing mutations may be required for tumorigenesis.

Interrogation of the mechanism through which the podocyte lineage might contribute to Wilms tumor and be further promoted by SIX1/2-Q177R will require an *in vitro* model system that recapitulates this developmental progression. Tran et al extended their findings to hiPSC-derived kidney organoid differentiations and illustrated that podocyte-like cells followed a developmental trajectory resembling that of *in vivo* hFK podocytes, characterized by a high degree of conservation of most of the identified EP and LP genes. Additionally, xenotransplantation of these avascular organoids in immunocompromised mice resulted in vascularization of the organoids by the host, as has been demonstrated similarly by several groups (Sharmin et al 2016, Van den Berg et al 2018, Ryan et al 2021, Koning et al 2022). Hence, hiPSC-derived kidney organoids represent an appropriate model system for future mechanistic studies of podocyte lineage contribution to tumorigenesis.

Our study and the findings herein are not without limitations. Wilms tumors are notoriously heterogenous and histological classification is fundamentally subjective. However, tumors classified as mixed histology imply the presence of some identifiable blastemal components, supporting our observation of the overall high degree of transcriptional correlation between tumors of different histological subtypes. Due to the limited availability of SIX1/2-Q177R tumor tissue, the nine samples included in this analysis undoubtedly fail to capture the full extent of variability among tumors with this mutation. Nonetheless, our use of stringent filtering, expression, and statistical thresholds increases the likelihood of biologically significant findings resulting from of our differential gene expression analysis. Conversely, these same stringent thresholds likely obscured some differentially expressed genes of biological significance. For example, NPC and podocyte marker *WT1*, as well as *CDH1* encoding E-cadherin, were excluded from this analysis due to several tumors not meeting the expression threshold. Additionally, the SIX1-Q177R ChIP-seq data utilized in this study was derived from a chemotherapy-treated tumor specimen (Wegert et al 2015). As such, this sample does not serve as a direct comparison to the SIX1-Q177R tumors used in the RNA-seq analysis. Therefore, inference of putative target gene regulation by SIX1-Q177R in chemotherapy-naïve tumors is only correlative at this time. Despite these limitations, our data support a reassessment of the differentiation status of Wilms tumors in which progression through podocyte lineage specification is a common feature of high-risk Wilms tumors of all histological subtypes, is further augmented in tumors harboring the relapse-associated SIX1/2-Q177R mutation, and is advantageous for oncogenesis in the developing hFK.

## ACKNOWLEDGEMENTS

The authors would like to thank Dr. Manfred Gessler for generously sharing the Wilms tumor ChIP-seq data, as well as the Therapeutically Applicable Research to Generate Effective Treatments (https://ocg.cancer.gov/programs/target) initiative (TARGET) study phs000218.v24.p8 for generating the Wilms tumor RNA-seq data used in this analysis (data available at https://portal.gdc.cancer.gov/projects). Work in this study was supported by the Maren Endowment and UNC Lineberger Tier 1 Pilot/Development Award (L.L.O.), NIH grant R01 HG010501 (M.L.B.), as well as NCI F31CA257443 and NIGMS Training Grant 5T32 GM007092 (M.J.S).

## AUTHOR CONTRIBUTIONS

L.L.O. conceptualized the study, interpreted data, and revised the manuscript. M.J.S. performed experiments, data analysis, interpreted data, and wrote the manuscript. U.P. performed luciferase enhancer assays and cloning related to luciferase enhancer assays. S.K.P. performed PBM experiments and data analysis. S.S.G. analyzed and interpreted PBM data. S.K.P. and S.S.G. contributed to the manuscript text. S.S.G. prepared PBM data figures. M.L.B. interpreted PBM data and supervised research.

## EXPERIMENTAL PROCEDURES

### Cell culture and transfections

HEK293T and MCF-7 (gift from Dr. Richard Cheney, UNC-Chapel Hill) cell lines were cultured in DMEM/F12 w/ L-glutamine and HEPES (Gibco) + 10% Fetal Bovine Serum (Omega Scientific) + 1X Penicillin/Streptomycin (Gibco), unless indicated elsewhere. Media was exchanged every 3-4 days and cells were passaged at confluence. Transfections for all assays were carried out using Lipofectamine 3000 kit (Invitrogen) following manufacturer’s protocol, scaled for tissue culture vessel used. Cells were assayed ~48 hours post-transfection.

### General cloning

All restriction digest reactions were carried out at 37C using CutSmart Buffer (New England Biolabs). All restriction enzymes were from New England Biolabs, unless indicated elsewhere. Restriction digest products were subjected to gel electrophoresis in 1% agarose gel + 1X SYBR Safe DNA Gel Stain (Invitrogen). DNA bands of digested DNA were excised and purified using NucleoSpin Gel and PCR Clean-Up kit (Machery-Nagel) following the manufacturer’s protocol. All ligations were carried out at RT using T4 DNA ligase (New England Biolabs) following the manufacturer’s protocol. Unless indicated elsewhere, plasmids were transformed in 5-alpha Competent *E. coli* cells (New England Biolabs) following the manufacturer’s protocol. LB agar plates containing 1X ampicillin (Sigma Aldrich) were then streaked with transformed bacteria and incubated at 37C overnight. Individual bacteria colonies were picked using P1000 pipet tips and cultured in LB broth + 1X ampicillin at 37C on orbital shaker from several hours to overnight. Plasmids were purified from cultures using NucleoSpin Plasmid EasyPure purification kit (Machery-Nagel) following manufacturer’s protocol. Plasmids were submitted to GENEWIZ (Azenta Life Sciences) for Sanger sequencing to verify DNA sequence. Purified plasmids verified by DNA sequencing were then either used directly in applicable assay or re-transformed in 5-alpha cells and an individual bacteria colony would be used to inoculate a 200mL liquid culture in LB broth + 1X ampicillin and culture would incubate overnight at 37C on orbital shaker. Plasmids were then purified using NucleoBond Xtra Midi Plus EF Kit (Machery-Nagel), following manufacturer’s protocol including use of NucleoBond finalizers.

### General SDS-PAGE and Western blots

All samples were prepared using LDS sample buffer (Invitrogen) + 2.5% 2-Mercaptoethanol (Sigma-Aldrich) and incubated at 70C for 10 minutes prior to electrophoresis. Samples were run on Novex 4-20% Tris-Glycine gels (Invitrogen) along with Precision Plus Protein All Blue Prestained Protein Standards (Biorad) using 1X Tris-Glycine running buffer (25mM Tris Base (Fisher BioReagents), 190mM Glycine (Fisher Chemical), and 3.5 mM SDS (Fisher Chemical)) in a Mini Gel Tank (Invitrogen). Proteins were then transferred to a nitrocellulose membrane (GE Water and Process Technologies) using a wet transfer protocol and a Mini Trans-Blot Cell (Bio-Rad) transfer apparatus at 4C, 100V for 1 hour in transfer buffer (192 mM Glycine, 25mM Tris Base, and 20% methanol). Membranes were then washed several times with ddH20. Membranes were blocked in PBS + 0.1% TWEEN-20 (Fisher BioReagents) + 5% dry milk at RT for 20-40 minutes and residual milk was washed from membrane with water. Primary antibody solutions were made in PBS + 0.1% TWEEN-20 + 3% BSA (Fisher BioReagents). Unless indicated elsewhere, membranes were incubated in primary antibody solution with gentle rocking at either RT for 1 hour or overnight at 4C. Membranes were washed at least three times with PBS + 0.1% TWEEN-20, 5 minutes per wash. Secondary antibody solutions were made in PBS + 0.1% TWEEN-20 + 1% dry milk. Membranes were incubated in secondary antibody solution for 1 hour at RT. Membranes were washed at least three times with PBS + 0.1% TWEEN-20 and then incubated in Pierce ECL Western Blotting Subtrate (Thermo Scientific) for 1-2 minutes at RT. Blots were imaged using the iBright FL1500 Imaging System (Invitrogen).

### ChIP-seq data analysis

SIX1 ChIP-seq data from Wilms tumors was kindly shared by Dr. Manfred Gessler (Wegert et al. 2015) in the form of bigWig files from two SIX1 wild type ChIP-seq samples and two SIX1-Q177R ChIP-seq samples (50 bp bins, normalized to pooled input by signal extraction scaling (SES)). BigWig files were converted to bedGraph format using UCSC tools bigWigToBedGraph (Kent et al 2010). Duplicates were merged and log_2_ fold enrichments over input were averaged between duplicates, generating a single bedGraph file for each set of duplicate samples. Genome coordinates were transformed from hg19 genome assembly to hg38 using the UCSC LiftOver tool (Kent et al 2002). Peaks were called using MACS2 bdgpeakcall (Zhang et al 2008) through the Galaxy web platform usegalaxy.org (Afgan et al 2016) using a log_2_ fold cutoff of +/− 2. Week 17 human fetal kidney SIX1 ChIPseq peak BED file was from O’Brien et al, 2016. Peak overlaps between samples were identified using bedtools (Quinlan and Hall 2010) intersect intervals in Galaxy. Peak sequences were obtained using bedtools GetFastaBed in Galaxy and these FASTA files were used as input for motif discovery in STREME (MEME Suite 5.4.1) (Bailey 2021) using default settings, or in SEA in MEME Suite 5.4.1 to assess enrichment of motifs of potential SIX1 co-factor MEIS1 (Bailey and Grant 2021 bioRxiv). The identified SIX1 or SIX1-Q177R motifs from STREME were also used as input into FIMO (Grant 2011) using p-value cutoffs of either <1E^-4^ or <1E^-5^. The peak coordinates containing the desired motifs were converted to BED format and duplicate coordinates within each file were removed using bedtools MergeBED in Galaxy. These BED files were then used as input in the Genomic Regions Enrichment of Annotations Tool (GREAT) (McLean et al 2010) using whole genome background and basal plus extension association rules, changing distal up to 500 Kb. Putative target genes were identified using a binomial false discovery rate cutoff of 0.2.

### Protein binding microarray

#### Cloning

DNA sequences flanked by XhoI and NdeI restriction sites and encoding for N-terminal GST-tagged-SIX1 or N-terminal GST-tagged-SIX1-Q177R homeodomains were synthesized by Integrative DNA Technologies (IDT) as gBlocks (Supplemental Table 2). PCR using Q5 High-Fidelity DNA Polymerase (New England Biolabs) was used to amplify gBlocks and PCR products were purified using Nucleospin Gel and PCR Cleanup kit following manufacturer’s protocol. DHFR control plasmid from PURExpress In Vitro Protein Synthesis Kit (New England Biolabs) was used as backbone for subsequent ligations. DHFR control plasmid and amplified gBlocks were digested with XhoI and NdeI restriction enzymes. Gel purification, ligation, transformation, and subsequent purification was performed as described in “Cloning” above.

#### In vitro transcription/translation and protein concentration quantification

*In vitro* transcription/translation was carried out using PURExpress In Vitro Protein Synthesis kit, following the manufacturer’s protocol. 1μL from each reaction was diluted 1:100 in nuclease-free water. 7μL of each dilution was then used in SDS-PAGE alongside a dilution series of recombinant GST protein (Sigma-Aldrich #SRP5348) from 5ng-200ng. SDS-PAGE and Western blot were performed as described above, with the following changes: 90 minute primary antibody incubation at RT (rabbit α-GST, Sigma Aldrich #G7781, 1:4000) and 30 minute secondary incubation at RT (goat α-rabbit HRP, 1:5000) (Supplemental Figure 1D). Protein concentration was quantified using band intensities obtained using the Gel tool in ImageJ (Schneider et al 2012). Standard curve was generated using known recombinant GST bands in gel. Using Microsoft Excel, a logarithmic line of best fit was generated and used to solve for mass of the *in vitro* transcribed/translated samples. Molarity of the purified protein samples was calculated using a molecular weight of 36.31 kDa, nuclease-free water was added to bring the molarity of each sample to 4.5μM. Aliquots were made and stored at −80C.

#### Protein binding microarrays

PBMs were performed on universal ‘all 10-mer’ arrays in 8 x 60K format (GSE AMADID # 030236, Agilent Technologies) essentially as described previously (Berger and Bulyk 2009, Berger et al 2006). PBM experiments were performed in duplicate at 300 nM final concentration of GST-tagged protein. Protein binding was detected with Alexa488-conjugated α-GST antibody (Invitrogen A-11001). Arrays were scanned using a GenePix 4400A microarray scanner (Molecular Devices). Raw data files were processed and binding was quantified using the Universal PBM Analysis Suite (Berger and Bulyk 2009). Motif position weight matrices were derived using the Seed-and-Wobble algorithm (Berger and Bulyk 2009, Berger et al 2006) and sequence logos were generated with enoLOGOS (Workman et al 2005). Pattern E-scores were generated using the same algorithm and input files as 8-mer E-scores (Berger and Bulyk 2009), with probes that contain matches to a given sequence pattern replacing probes containing a given 8-mer as the foreground in the calculation.

### Wilms tumor RNA-seq data analysis

Wilms tumor RNAseq data was obtained through the National Cancer Institute TARGET Data Matrix (https://ocg.cancer.gov/programs/target/data-matrix) in the form of gene quantification text files. From each gene quantification file, raw counts were transferred to an excel spreadsheet to create a count matrix. This count matrix was then used in Galaxy for differential gene expression analysis using limma-voom with sample quality weights, filtering out lowly expressed genes with CPM < 2 if threshold was not met in all samples, and using a log_2_ fold change cutoff of +/− 1.5, and all other default settings (Smyth 2005, Law et al 2014, Liu et al 2015). A matrix of Pearson’s correlation coefficients was generated between all tumor samples using all genes that passed limma-voom low expression filtering, excluding duplicate genes. Log_2_ CPM values were first scaled using the scale() function in R and correlation coefficients were generated using the cor() function. Unsupervised hierarchical clustering and heatmap generation was performed using the pheatmap package with default clustering parameters (Kolde, 2019). Violin plots shown in article figures and supplemental figures were generated using the ggplot2 package in R (Wickham, 2016).

### Single cell RNA-seq analysis

Raw and processed data was obtained from three studies: GSE112570, GSE139280, GSE124472 (only sample GSM3534656). All subsequent data processing and analyses was performed using the Seurat package and following the analysis workflows outlined in vignettes “PBMC 3K guided tutorial” and “Introduction to scRNA-seq integration” (https://satijalab.org/seurat/index.html, Hao and Hao et al 2021, Stuart and Butler et al 2019, Butler et al 2018, Satija and Farrell et al 2015). Briefly, each dataset was filtered using the following parameters: LindströmWk14 – nFeature_RNA = 1500-4000, mitochondrial counts < 5%; LindströmWk17 – nFeature_RNA = 1000-3500, mitochondrial counts < 5%; TranWk17zone1 – nFeature_RNA = 1000-5000, mitochondrial counts < 5%. Each dataset was normalized independently and variable features were identified independently. Integration features were selected and integration anchors were identified. An integrated assay was then created, data was scaled, PCA and UMAP dimensional reduction were performed using n=20 principle components/dimensions. Neighbors were found and clusters were found using resolution = 0.5. Cluster markers were found using FindAllMarkers() function, min.pct = 0.15, logfc.threshold = 0.25. AverageExpression() function was used to calculate average expression value for each cluster, used return.seurat = TRUE to return SeuratObject with scaled and centered expression values generated from ScaleData() function. Dot plots were generated using DotPlot() function, heatmaps were generated using DoHeatmap() function, and violin plots were generated using VlnPlot() function. Ridgeline plots were generated using the ggridges package in R (Wilke, 2022).

### Luciferase assays

#### Cloning and plasmids

pBV-Luc was a gift from Bert Vogelstein (Addgene plasmid # 16539; http://n2t.net/addgene:16539; RRID:Addgene_16539). This plasmid encodes firefly luciferase driven by a minimal promoter element and was digested with NheI and HindIII, however this digestion removed the minimal promoter from the pBV-Luc vector. SIX1_enhancer gBlocks (Supplemental Table 2) were PCR amplified and digested with NheI and HindIII restriction enzymes and annealed with digested pBV-Luc. To re-insert the minimal promoter sequence, single-stranded DNA oligos containing the minimal promoter sequence flanked by 5’-HindIII and 3’-NcoI restriction sites (Supplemental Table 2) were annealed in 1X annealing buffer (10mM Tris Base, 50mM NaCl (Fisher Chemical), 1mM EDTA (Invitrogen)) and incubated on thermocycler at 95C for 2 minutes followed by cooling to 25C at a rate of −0.1C/second. Annealed minimal promoter oligo and SIX1_enhancer-pBV-Luc were then digested with HindIII and NcoI restriction enzymes and annealed. This plasmid was then used for all subsequent cloning of WNT5A proximal and distal CRE luciferase constructs using NheI/HindIII restriction sites (Supplemental Table 2).

pRL-SV40P was a gift from Ron Prywes (Addgene plasmid # 27163; http://n2t.net/addgene:27163; RRID:Addgene_27163) and was used as renilla luciferase expression control. Empty pCIG and empty pCS2+ plasmids were used as empty vectors for total DNA transfection normalization. EYA1-2xHA fragment was generated by PCR amplifying EYA1 coding sequence from pCS2+-EYA1-FLAG plasmid, swapping out FLAG tag for 2xHA tag (Supplemental Table 2). pCS2+-EYA1-2xHA plasmid was generated using EcoRI and XbaI restriction sites in EYA1-2xHA fragment and pCS2+ plasmid. pCIG-SIX1 and pCIG-SIX1-Q177R plasmids were generated using gBlocks synthesized by Integrated DNA Technologies (IDT) containing the coding sequences for the respective proteins (Supplemental Table 2). The gBlocks were PCR amplified and flanking ClaI and XhoI restriction sites were added and used for subsequent digestion and ligation into pCIG vector.

#### Luciferase assay

MCF-7 cells were cultured in 6-well plates to ~90% confluence and media was exchanged prior to transfection. For each biological replicate of each DNA element assayed, three wells would be transfected with 5 μg total DNA/well: one no protein control condition, one SIX1/EYA1 condition, and one SIX1-Q177R/EYA1 condition. Control transfections consisted of 500 ng firefly luciferase vector, 10 ng renilla luciferase vector, 1.5 μg empty pCIG, and 3 μg empty pCS2+. Luciferase assays were performed using the Dual Luciferase Reporter Assay System (Promega). Each well was harvested using a cell scraper and 100 μL 1X Passive Lysis Buffer. After trituration with a P200 pipet tip, 20 μL lysate was transferred to each of 3x wells of a clear, flat-bottom 96-well plate for technical triplicate per condition. Luminescence was measured using a BioTek Synergy HT plate reader with the following settings: 10 second integration time, 135 gain, 1 mm read height. 100 μL/well Luciferase Assay Reagent II was dispensed into all wells containing cell lysate using P200 multichannel pipet and plate was immediately placed in plate reader. After firefly luciferase luminescence was measured, 100 μL/well Stop & Glo reagent was dispensed using P200 multichannel pipet and plate was immediately placed in plate reader. Fold change relative to no protein control was calculated by comparing ratios of firefly/renilla luminescence from SIX1 or SIX1-Q177R conditions to that of no protein control.

### Protein expression and purification

The same SIX1 and SIX1-Q177R gBlocks used for pCIG cloning were PCR amplified to add flanking BamHI and XhoI restriction sites. Digested gBlocks were ligated to digested pGEX-6p1-N-HA (gift from Andrew Jackson & Martin Reijns, Addgene plasmid # 119756; http://n2t.net/addgene:119756; RRID:Addgene_119756). BL21 (DE3) Competent *E. coli* (New England Biolabs) were transformed following manufacturer’s protocol and streaked LB agar + 1X ampicillin plates were incubated overnight at 37C. One colony was picked from each transformation and cultured in 14 mL tubes each containing 7 mL LB broth + 1X ampicillin in orbital shaker at 37C overnight. Glycerol stocks were made from each overnight liquid culture by mixing 50% glycerol solution and liquid bacteria culture at a 1:1 ratio, then stored at −80C. A pipet tip was used to transfer a small amount of each glycerol stock to flasks containing 50 mL LB broth + 1X ampicillin and cultures were incubated on orbital shaker overnight at 37C. Each 50 mL culture was transferred to a 2 L flask containing 950 mL LB broth + 1X ampicillin and cultured at 37C on orbital shaker until OD600 = 0.55-0.56. 5 mL of 100mM IPTG (Sigma-Aldrich) solution was added to each culture and flasks were incubated on an orbital shaker at 25C for 19 hours. For each culture, the entire culture volume was distributed into four 250 mL centrifuge bottles and centrifuged at 5,500 RPM for 20 minutes at 4C. All pellets for each culture were resuspended and pooled in 35 mL supernatant and the final suspension was transferred to 50 mL tubes. Tubes were centrifuged at 4,000 x g for 18 minutes at 4C. Supernatants were discarded and pellets stored at −80C.

Frozen bacteria pellets were thawed on ice and loosened in 25 mL lysis buffer (20 mM Tris-HCl, 150 mM NaCl, 1% Triton X100, 10 mM DTT, 1X Protease Inhibitor Cocktail (Roche)). Samples were sonicated on ice in a 4C cold room using a Branson Sonifier 250, 10 cycles of 10 sec ON/OFF at 50% amplitude/duty cycle. Tubes were incubated on ice for 15 minutes then centrifuged at 13,000 x g for 15 minutes at 4C. Water was decanted from a glutathione-agarose bead mixture (0.84 g. glutathione-agarose beads in 168 mL water, incubated at 4C overnight). 120 mL lysis buffer was added to the beads, mixed, and incubated at 4C for 20 minutes. Lysis buffer was decanted from the beads and 8 mL of bead slurry transferred to each of 2x 50 mL tubes. Bacterial supernatants were added to tubes containing beads and tubes were put on a tube rotator at 4C for 1.5 hours. EconoColumns (BioRad) were wet and washed with 1x column volume PBS + 1% Triton-X100 and then emptied. Supernatant/bead mixtures were added to columns and column spigots opened full to allow gravity flow. As flow was near stopping, beads were washed with full column volumes of cleavage buffer (50 mM Tris-HCl, 1 mM EDTA, 1 mM DTT, 1% Triton-X100 (Fisher BioReagents)) four times. 200 μL of PreScission Protease (Cytiva) was mixed with 9.8 mL cleavage buffer, 5 mL added to each column, inverted to mix, and incubated at 4C for 2 hours, inverted to mix, and incubated at 4C for an additional 2 hours. Flow-through was collected in 15 mL conical tubes. 12-14,000 MWCO dialysis tubing (Spectrum Laboratories) was incubated at RT in 1.5L H2O + 5 mM EDTA for 2-3 hours, then rinsed thoroughly with H2O.

The entire volume of flow through was transferred to dialysis tubing, ends were clipped shut and incubated overnight at 4C submerged in 1 L dialysis buffer (50 mM Tris-HCl, 1 mM EDTA, 0.8 mM DTT) with gentle stirring. Used dialysis buffer was discarded, replaced with fresh 1 L dialysis buffer, and incubation continued at 4C for 2.5 hours. Contents of each dialysis tubing were transferred to 15 mL conical tubes. Membranes of 4x Amicon Ultra-4 30,000 MWCO Centrifugal Filter Units (Millipore) were pre-washed with water, then removed. For each protein solution, 2.5 mL was transferred to each of 2x filter units. Tubes were centrifuged at 3,000 x g for 25 minutes at 4C. For each protein concentrate, volumes were pooled from filter units. Protein concentration was measured using Pierce BCA Assay Kit (Thermo Scientrific) and following manufacturer’s protocol with the following changes: in 96-well plate BSA controls were loaded and 10 μL of a 1:10 sample dilution with 190 μL working reagent in duplicate were incubated at 37C for 30 minutes. Absorbance was measured at 562 nm on BioTek Synergy HT plate reader. BSA controls were used to generate a standard curve and the protein concentration of each purified protein sample was calculated. Dialysis buffer was added to each protein solution to bring concentrations to 2 mg/mL; aliquots were stored at −80C. Protein purification was validated by SDS-PAGE of a dilution series of each protein solution, followed by Western blot using α-SIX1 antibody (Cell Signaling Technology #12891) (Supplemental Figure 4B).

### Electrophoretic Mobility Shift Assays

Single-stranded DNA oligonucleotide probes were synthesized by IDT and biotin end-labeled using the Pierce 3’ Biotin end-labeling DNA kit (Thermo Scientific) following manufacturer’s protocol with the following changes: 25 pmol oligo per reaction were labeled, reactions stopped with 1 μL 0.5M EDTA, and complementary oligo labeling reactions were mixed prior to centrifugation at 13,000 x g for 2 minutes. For unlabeled oligos, 50 μL H2O was mixed with 25 μL of each complementary 100 μM oligo. Annealing buffer was added to 1X and annealed following the same procedure as described for the luciferase assay; annealed oligos were stored at −20C.

Purified SIX1 and SIX1-Q177R protein was diluted in water to the following molarities (nM): 1, 2, 4, 10, 20, 40, 100, 150, 200, and 300. EMSAs were performed using either Gelshift Chemiluminescent EMSA Kit (Active Motif) or LightShift Chemiluminescent EMSA Kit (Thermo Scientific) following manufacturer’s protocol for setting up binding reactions, with the following changes: glycerol and poly d(I-C) were not included in reactions, 3 μL/reaction of 1:10 diluted biotin end-labeled probe was used, 12 μL/reaction water was used (13μL for no protein control reaction), 1μL of the appropriate protein dilution per reaction was added and reactions incubated at RT for 25 minutes. A 6% DNA Retardation Gel (Invitrogen) was pre-ran at 100V for 30 minutes in 0.5X TBE buffer (45 mM Tris Base, 45 mM boric acid, 1 mM EDTA) in a Mini Gel Tank. 5 μL/reaction 5X Loading Dye was added, 20 μL/reaction loaded in the gel, and the gel ran at 100V for one hour. The transfer used a wet transfer protocol similar to that used for Western blot described previously but using 0.5X TBE buffer as transfer buffer and transferring to Immobilin Ny+ nylon membrane (Millipore). Membrane was put on paper towel to dry briefly. Membrane was placed face-down in a BioDoc-IT gel imager (UVP), the UV lamp turned on, and membrane incubated for 15 minutes. Crosslinked membrane was then either stored at −20C or proceeded directly to staining following kit manufacturer’s protocol. Stained membranes were imaged using iBright FL1500 Imaging System. ImageJ (Schneider et al 2012) was used to obtain intensities of unbound and bound probe bands. Percent probe bound was calculated as follows: (bound probe intensity / (bound probe intensity + unbound probe intensity)) x 100.

### Co-immunoprecipitation and Western Blot

The pCS2+-MEIS1-3xFLAG construct was generated by PCR amplification of *MEIS1* from human induced pluripotent stem cell-derived cDNA, followed by ligation into digested pCS2+ plasmid using BamHI and XhoI restriction sites (Supplemental Table 2). HEK293 cells were grown in 6 cm dishes and transfected as described above, but using 4 μg total DNA per dish, 2 μg/plasmid. One dish was transfected with only MEIS1-3xFLAG, one dish with MEIS1-3xFLAG and pCIG-SIX1, one dish with MEIS1-3xFLAG and pCIG-SIX1-Q177R, and one dish with only pCIG-SIX1 (Supplemental Table 2). Nuclear protein lysate was extracted ~48 hours post-transfection using Active Motif Nuclear Complex Co-IP kit following manufacturer’s protocol for preparation of nuclear extract, with the following changes: doubled digestion buffer volume used for each sample and incubated in digestion buffer at 37C for 25 minutes. 1 μL/sample was saved as input and 75 μL/sample used for immunoprecipitations. For each sample, 50 μL Protein G Dynabeads (Invitrogen) were washed 3x with 1 mL/wash PBS + 0.1% Triton X-100 + 0.5 mg/mL BSA. Beads were resuspended in 200 μL PBS + 0.1% Triton X-100 + 0.5 mg/mL BSA. 5 μL α-SIX1 antibody was added (Cell Signaling Technology, SIX1 (D4A8K) Rabbit mAb #12891), then incubated with gentle rotation at 4C for at least one hour. Beads were washed 3x with 1 mL/wash PBS + 0.1% Triton X-100 + 0.5 mg/mL BSA, final wash was removed. 200 μL PBS + 0.1% Triton X-100 + 0.5 mg/mL BSA was added to each tube along with 75 μL/tube of appropriate nuclear lysate. Beads/lysate mixtures were incubated with gentle rotation at 4C for one hour. Beads were washed 5x with 1 mL/wash PBS + 0.25% Triton X-100. The final wash was removed and 30 μL/tube LDS sample buffer (Invitrogen) + 2.5% 2-Mercaptoethanol (Sigma-Aldrich) was added to each immunoprecipitation tube and each input tube. Tubes were then incubated at 70C for 10 minutes. Samples were run on a 4-20% Tris-Glycine Novex gel and transferred to nitrocellulose membrane similar to as detailed above but transferred at 30V at 4C overnight. The membrane was washed and incubated with primary and secondary antibodies following the procedure described above but using the following antibodies: mouse α-FLAG (Sigma #F3165, 1:500) and donkey α-mouse HRP (Invitrogen #A16017, 1:10,000).

## Notes

### Competing Interest Statement

The authors have declared no competing interest.

### Summary of Updates

Protein binding microarray data has been added, authors have been updated to reflect the collaborative contributions

