## Supplemental data for "Podocyte lineage marker expression is preserved across Wilms tumor subtypes and enhanced in tumors harboring the SIX1/2 p.Q177R mutation"

**Supplemental Figure 1A**

SIX1 tumor only peaks motif discovery

| Motif | Logo | p-value | E-value | Sites | Distribution |
|---|---|---|---|---|---|
| 1-GWAACYTGAK | 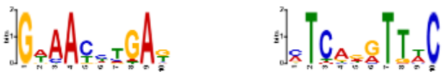 | 4.7e-071 | 2.1e-069 | 6932 (71.0%) | 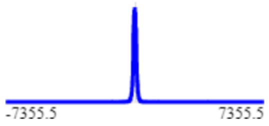 |

SIX1-Q177R only peaks motif discovery

| Motif | Logo | p-value | E-value | Sites | Distribution |
|---|---|---|---|---|---|
| 1-AAYHHRACACC | 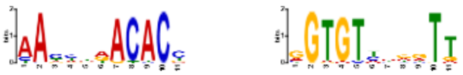 | 1.7e-040 | 7.6e-039 | 3944 (41.6%) | 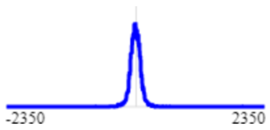 |

Shared Tumor peaks motif discovery

| Motif | Logo | p-value | E-value | Sites | Distribution |
|---|---|---|---|---|---|
| 1-GWAAYYTGABMC | 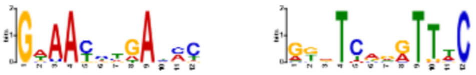 | 5.6e-072 | 3.2e-070 | 5999 (75.1%) | 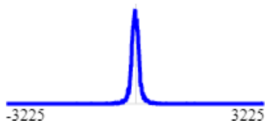 |

Wk17hFK SIX1 ChIP-seq motif discovery

| Motif | Logo | p-value | E-value | Sites | Distribution |
|---|---|---|---|---|---|
| 1-GWAAYYTGAKMYB | 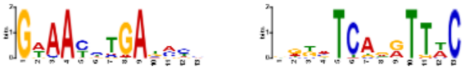 | 1.3e-110 | 6.6e-109 | 6178 (64.2%) | 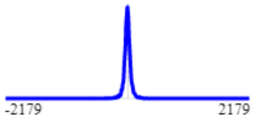 |

**Supplemental Figure 1B**

Wk17hFK peaks – MEIS1 motif enrichment

| Logo | Database | ID | Alt ID | *P*-value | *E*-value | Q-value | TP | FP | Enrichment Ratio | Score Threshold | Positional Distribution |
|---|---|---|---|---|---|---|---|---|---|---|---|
| 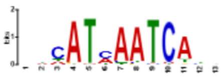 | MA1639.1 | MA1639.1 | MEIS1 | 1.47e-15 | 1.47e-15 | 1.47e-15 | 973 / 8666 (11.2%) | 655 / 8666 (7.6%) | 1.48 | 9.73 | 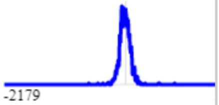 |

Shared Tumor peaks – MEIS1 motif enrichment

| Logo | Database | ID | Alt ID | *P*-value | *E*-value | Q-value | TP | FP | Enrichment Ratio | Score Threshold | Positional Distribution |
|---|---|---|---|---|---|---|---|---|---|---|---|
| 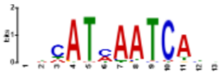 | MA1639.1 | MA1639.1 | MEIS1 | 4.11e-14 | 4.11e-14 | 4.11e-14 | 251 / 7186 (3.5%) | 110 / 7186 (1.5%) | 2.27 | 13.42 | 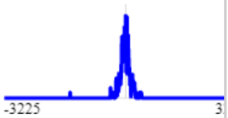 |

SIX1 tumor only peaks – MEIS1 motif enrichment

| Logo | Database | ID | Alt ID | *P*-value | *E*-value | Q-value | TP | FP | Enrichment Ratio | Score Threshold | Positional Distribution |
|---|---|---|---|---|---|---|---|---|---|---|---|
| 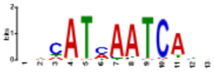 | MA1639.1 | MA1639.1 | MEIS1 | 1.61e-2 | 1.61e-2 | 1.61e-2 | 219 / 8784 (2.5%) | 174 / 8784 (2.0%) | 1.26 | 12.57 | 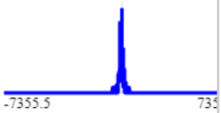 |

Q177R tumor only peaks – MEIS1 motif enrichment

| Logo | Database | ID | Alt ID | *P*-value | *E*-value | Q-value | TP | FP | Enrichment Ratio | Score Threshold | Positional Distribution |
|---|---|---|---|---|---|---|---|---|---|---|---|
| 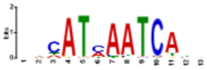 | MA1639.1 | MA1639.1 | MEIS1 | 2.23e-40 | 2.23e-40 | 2.23e-40 | 1018 / 8535 (11.9%) | 505 / 8535 (5.9%) | 2.01 | 10.47 | 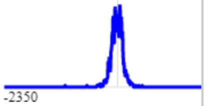 |

**Supplemental Figure 1C**

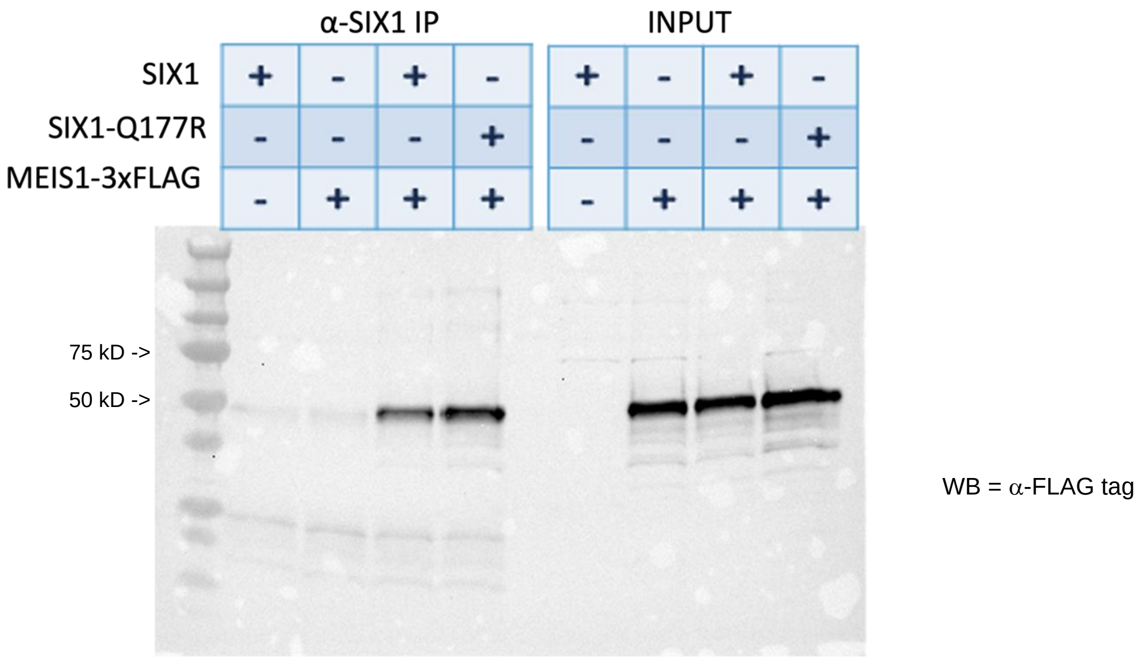


**Supplemental Figure 1D**

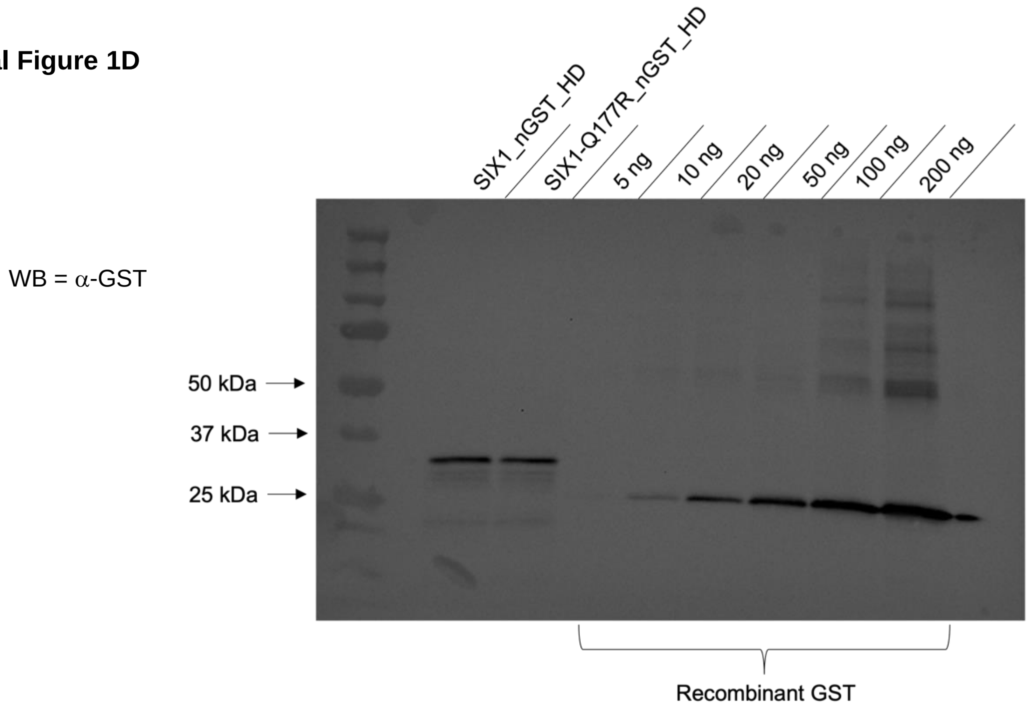


**Supplemental Figure 1: A)** Motif discovery output from STREME tool. Peak sequences used had been derived from after a first round of motif discovery using STREME followed by use of the FIMO tool to extract sequences containing SIX1-like motif. **B)** Motif enrichment output from SEA tool using same peak sequences used in panel A and searching for enrichment of MEIS1 DNA binding motif (JASPAR #MA1639.1). **C)** Western Blot using α-FLAG tag antibody following SIX1 immunoprecipitation and SDS-PAGE. **D)** Western Blot using α-GST antibody that was used to quantify concentration of *in vitro* transcribed/translated SIX1_nGST_HD and SIX1-Q177R_nGST_HD protein fragments alongside dilution series of recombinant GST protein.

**Supplemental Figure 2**

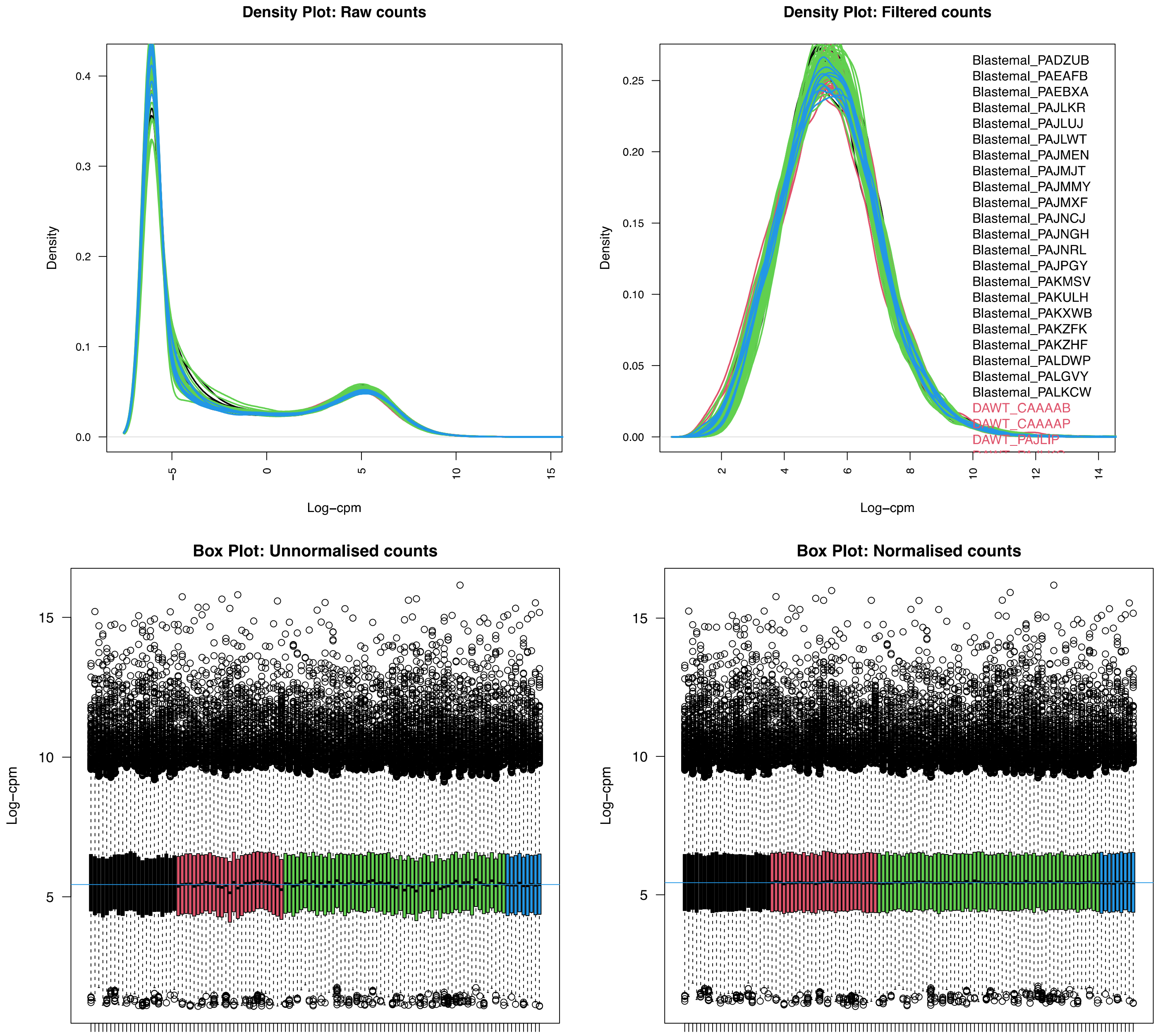

#### Supplemental Figure 2 continued

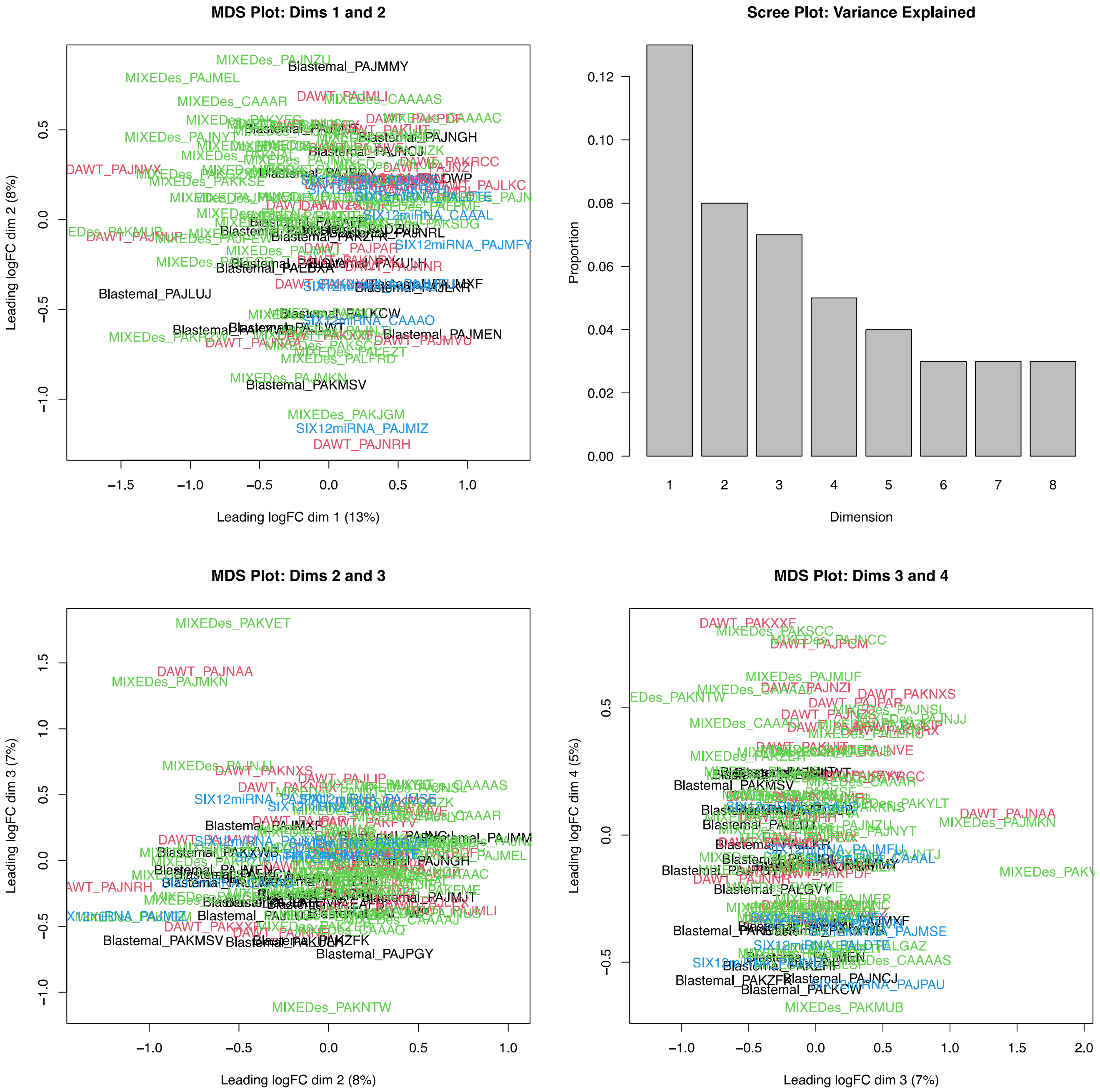

**Supplemental Figure 2 continued**

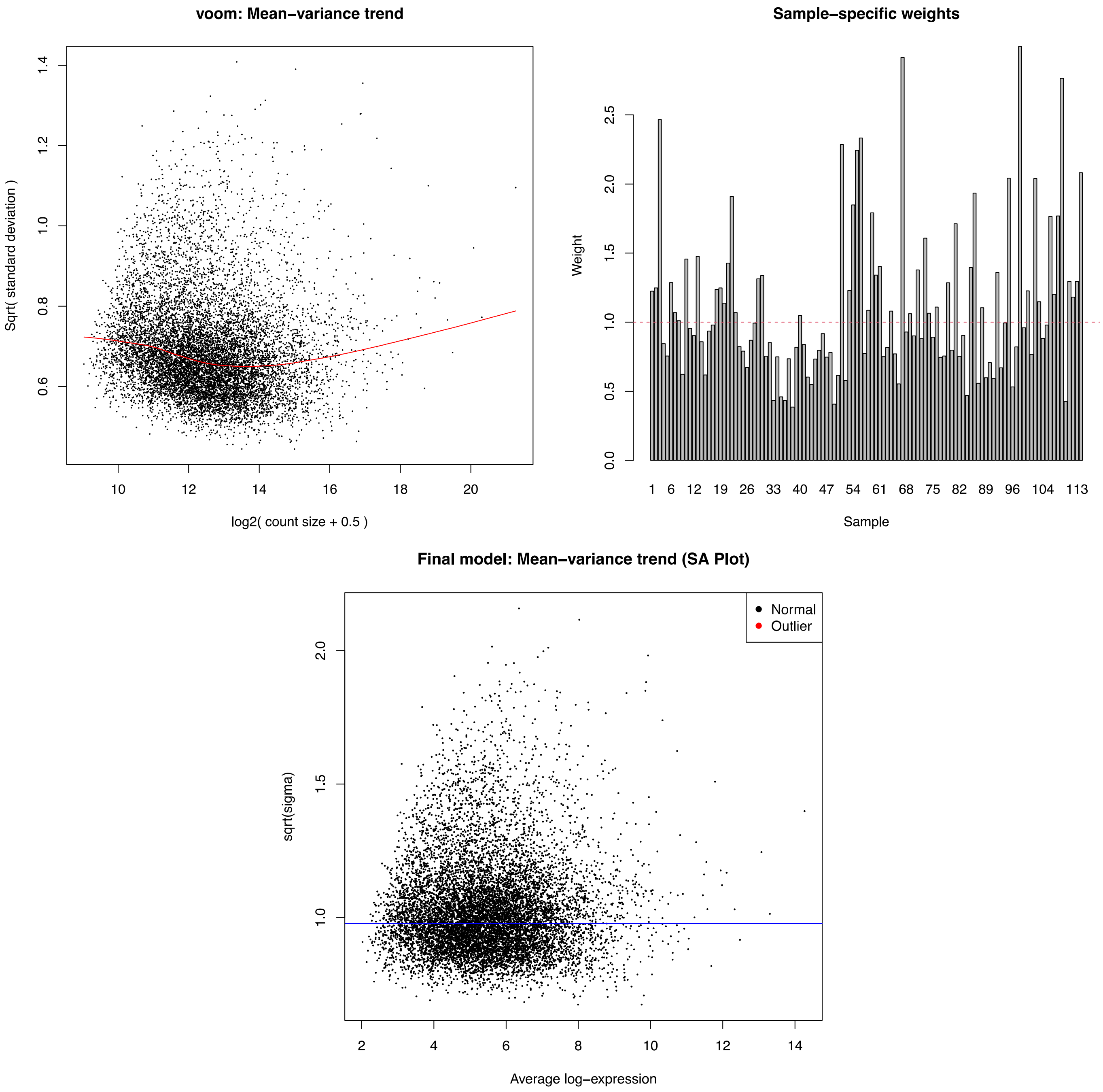


**Supplemental Figure 2:** Various plots/charts of metrics from limma-voom differential gene expression analysis.

### Supplemental Figure 3A

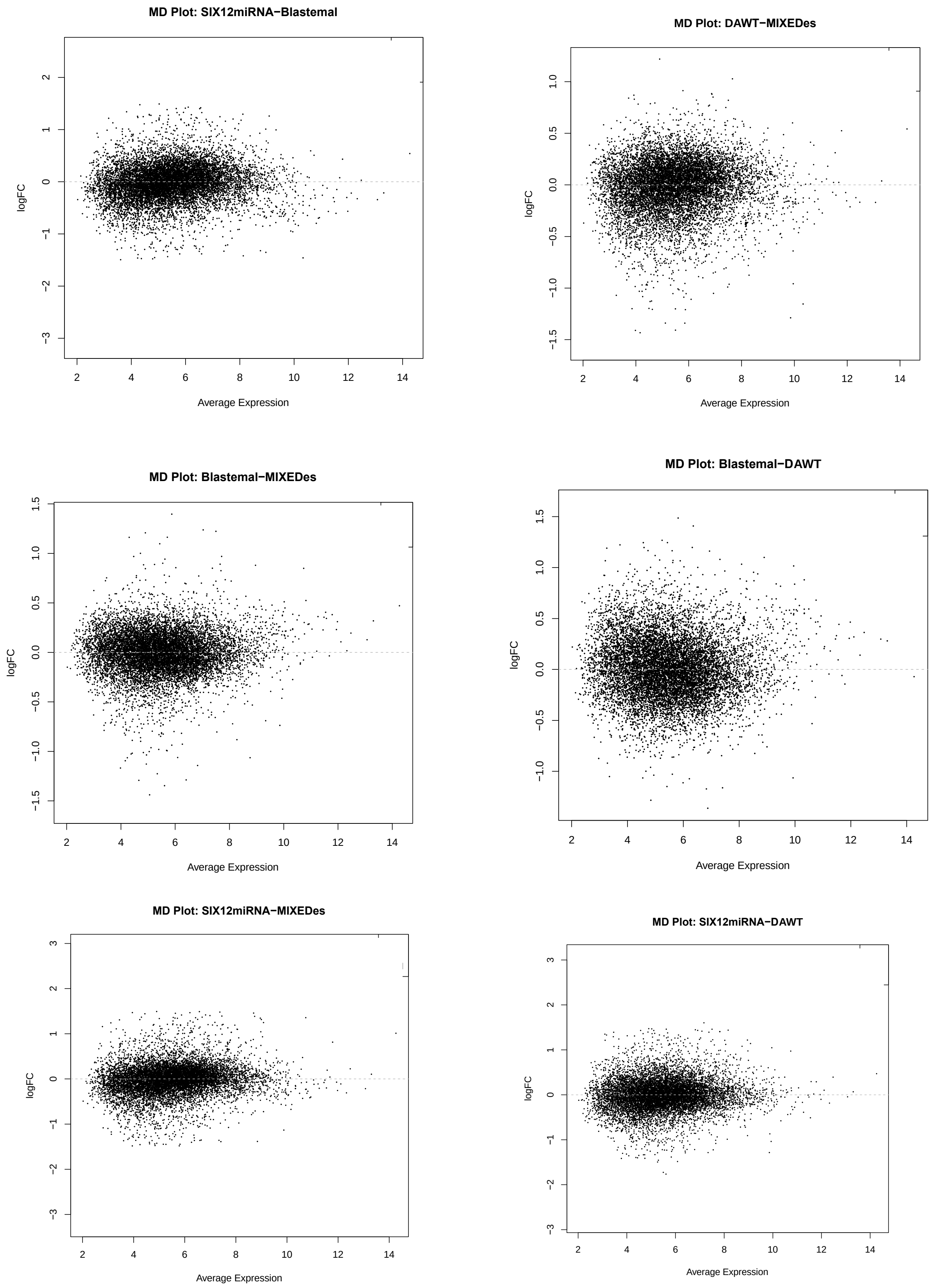

**Supplemental Figure 3B**

Downregulated in SIX1/2miRNA vs Blastemal (log2 FC > |1.5|, adj p < 0.05)

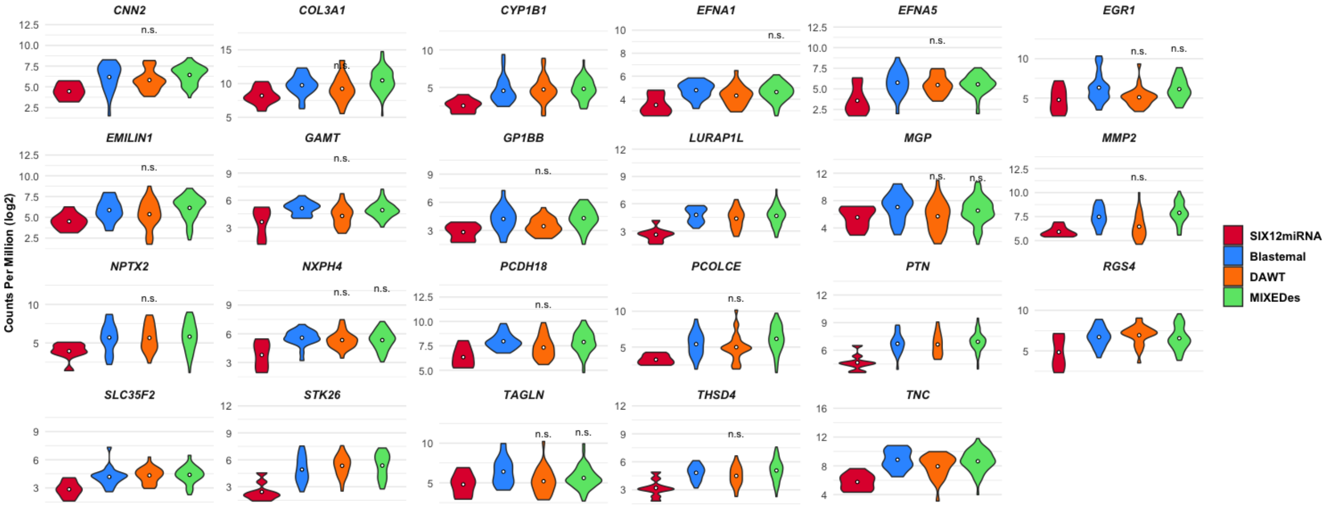


**Supplemental Figure 3: A)** MD plots from results of limma-voom differential gene expression analysis. Red dots indicate genes with > 1.5 log2 fold change and adj p-value < 0.05, blue dots indicate genes with < -1.5 log2 fold change and adj p-value < 0.05, Average Expression = log2 CPM. **B)** Violin plots showing the distributions of log2 counts per million (CPM) of the indicated gene set across tumor groups. Dot within each group plot represents the mean. Unless indicated by n.s. (not significant), the log2 fold change of that gene in the SIX1/2miRNA group was > |1.5| and adjusted p-value < 0.05 with respect to that tumor group.

**Supplementary Figure 4A**

***KDM2B***

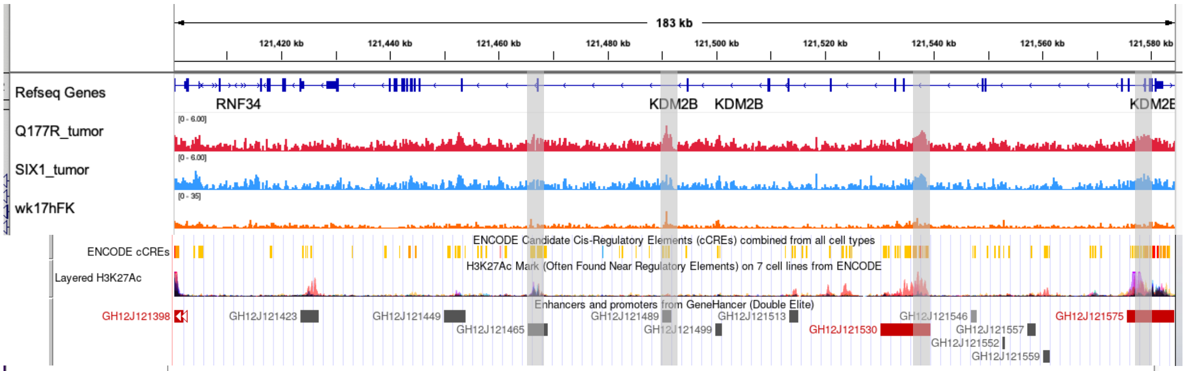


***CDKN1C***

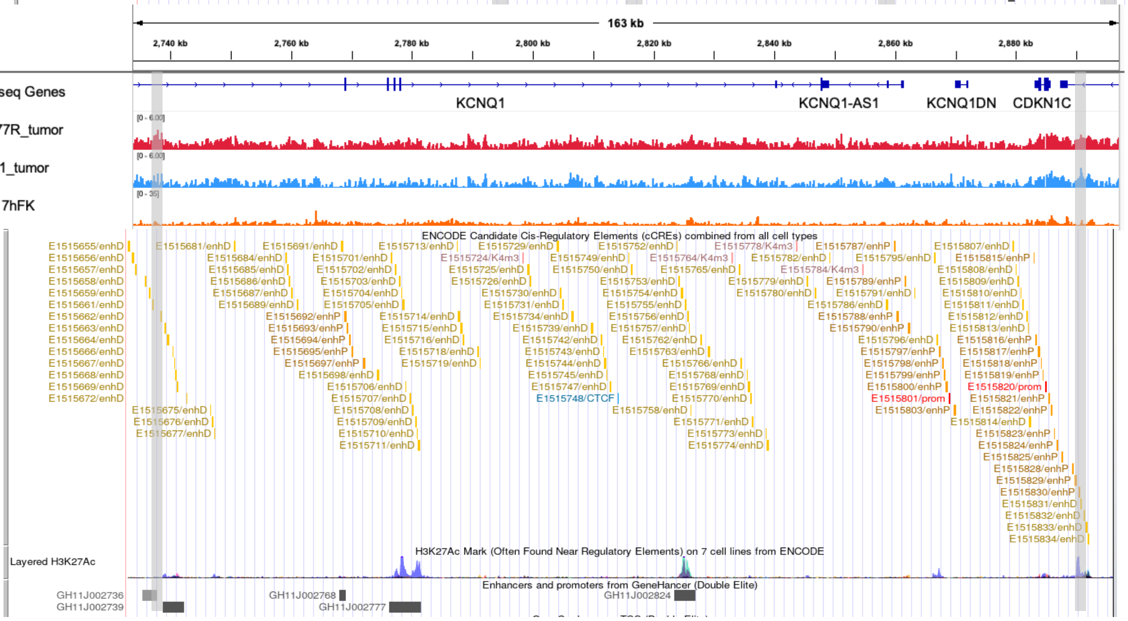


***FOXC1***

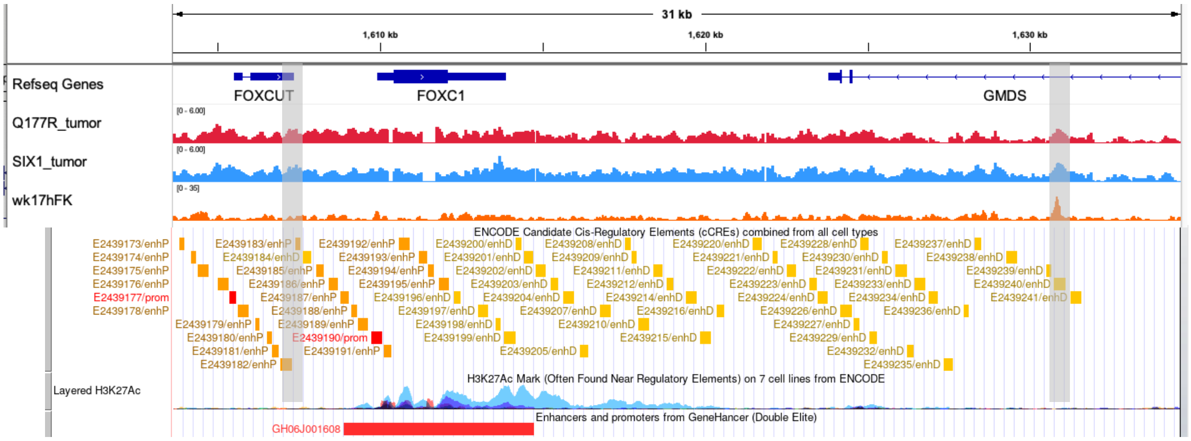

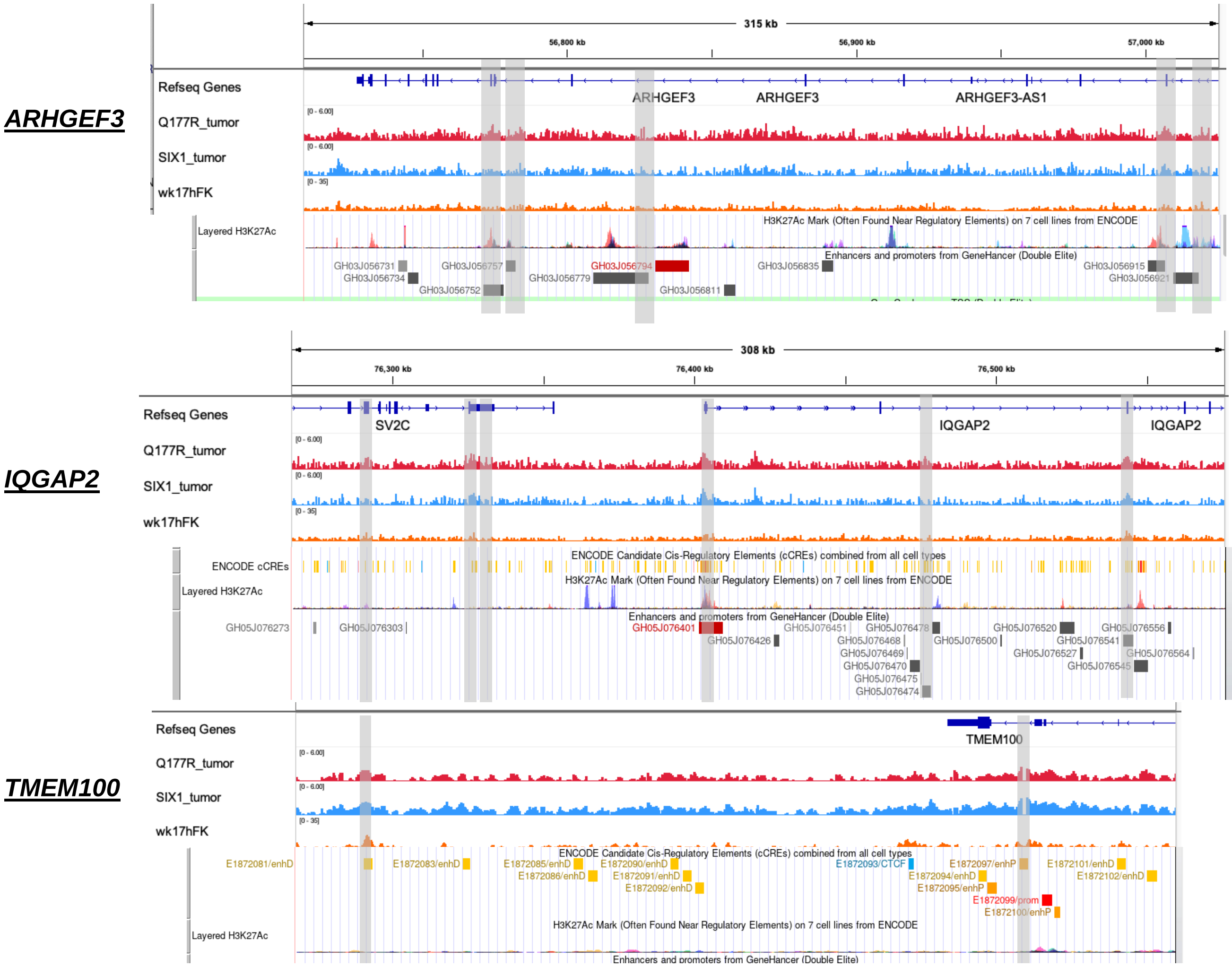


**Supplemental Figure 4B**

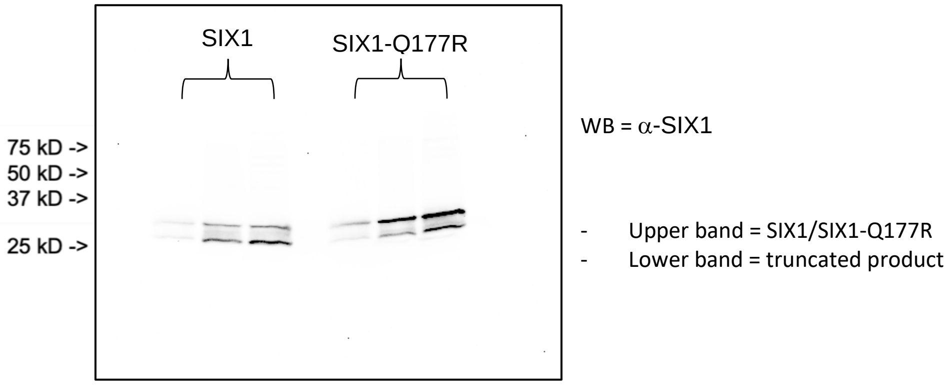


**Supplemental Figure 4:** **A)** IGV browser snapshots showing Q177R tumor, SIX1 tumor, and wk17hFK SIX1 ChIP-seq peak tracks, grey-shaded bars indicate locations of SIX1-Q177R called peaks. Below IGV browser snapshots are corresponding UCSC genome browser snapshots showing ENCODE candidate cis-regulatory elements, layered H3K27Ac, and GeneHancer predicted enhancer and promoter tracks. **B)** Western blot using α-SIX1 antibody following SDS-PAGE of increasing concentrations of purified SIX1 and SIX1-Q177R.

**Supplemental Figure 5A**

**Lindstrom Wk 14 kidney 1**

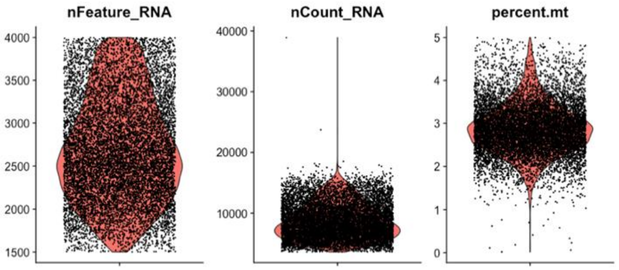


**Lindtsrom Wk 14 kidney 2**

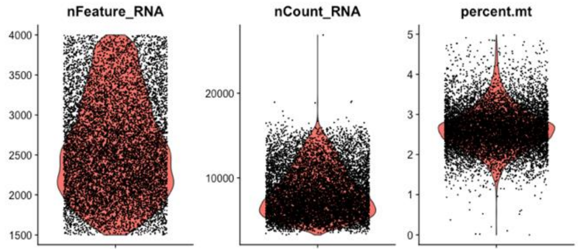


**Lindstrom Wk 17 kidney 1**

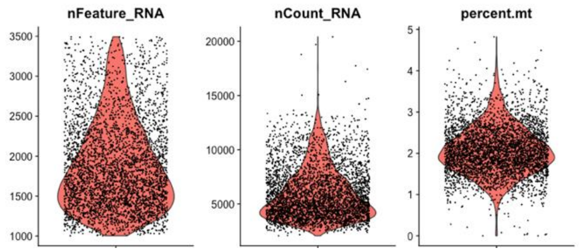


**Lindtsrom Wk 17 kidney 2**

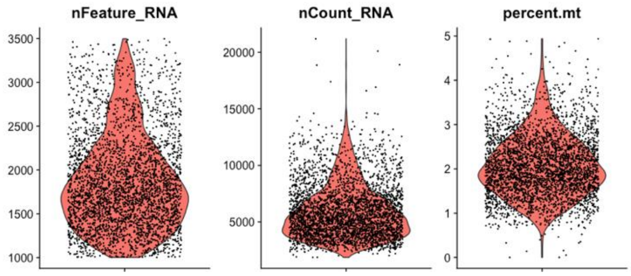


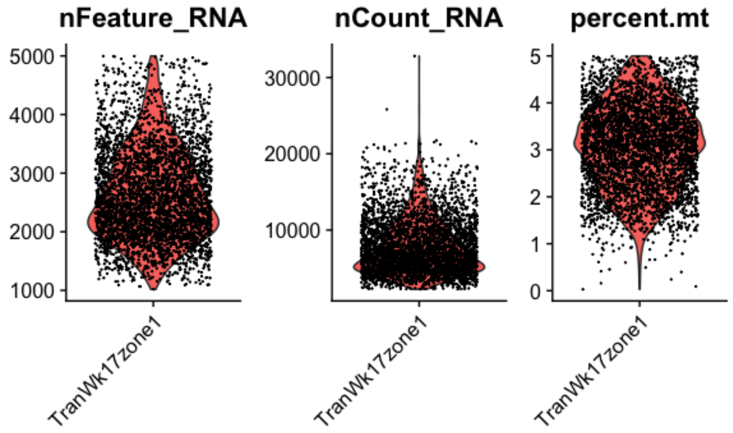

**Supplemental Figure 5B**

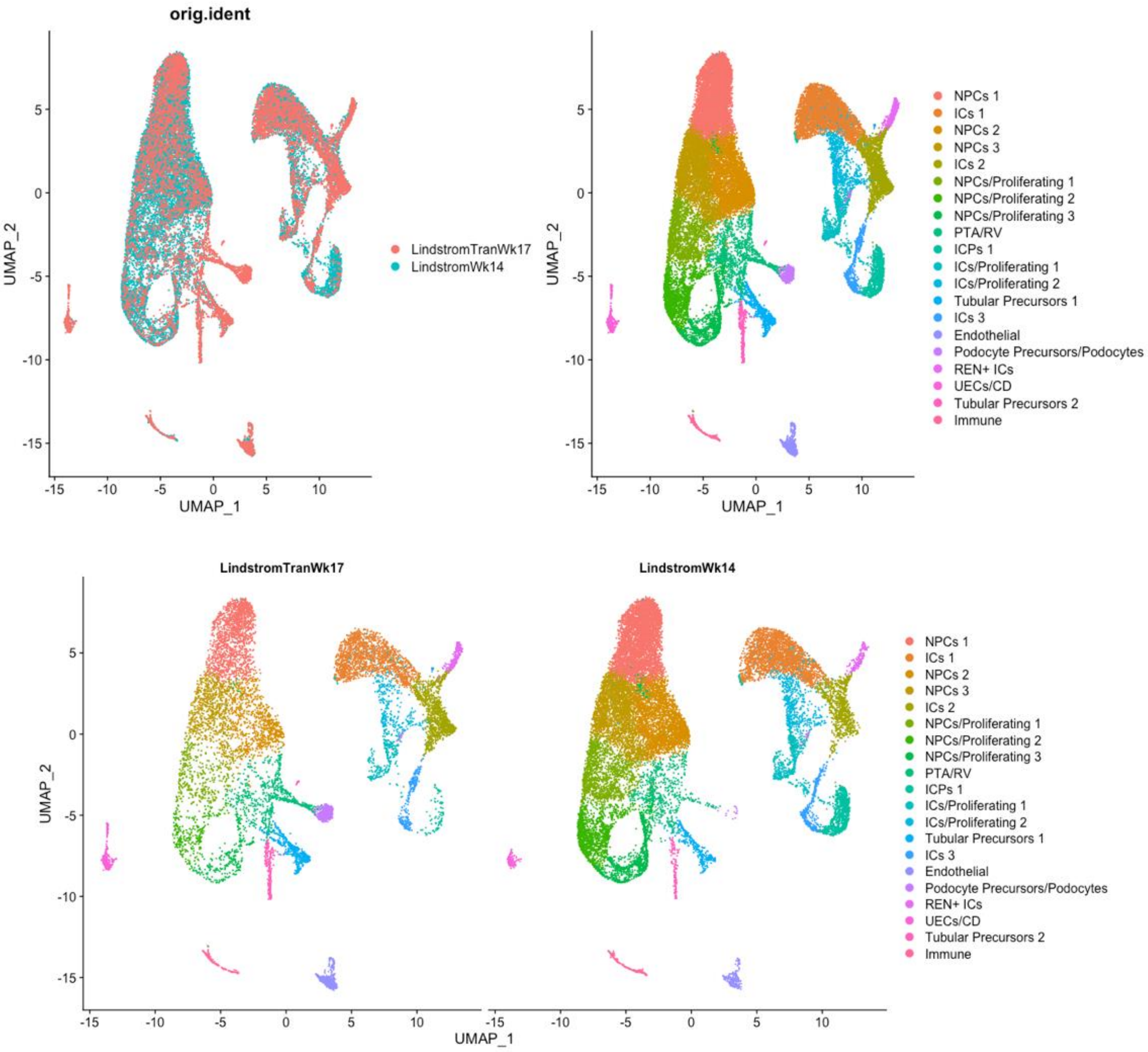


**Supplementary Figure 5C**

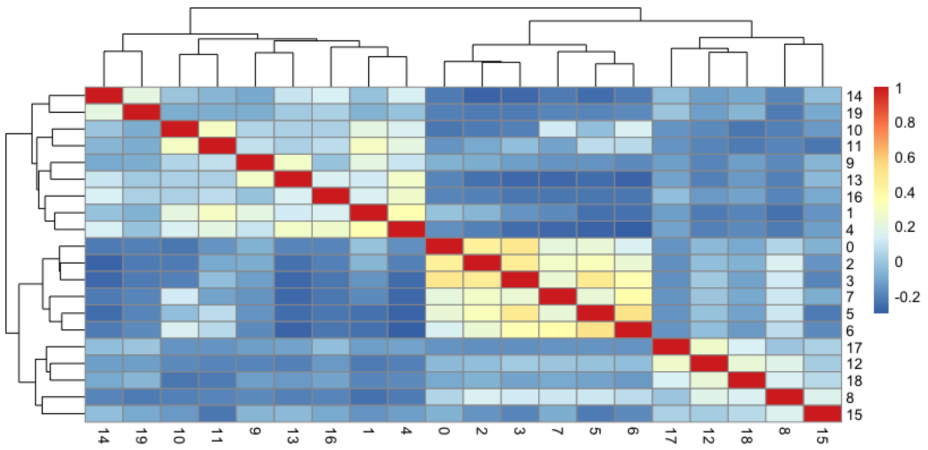


**Supplementary Figure 5: A)** Plots of QC metrics for cells in each of the indicated single cell RNA-seq datasets after data processing. **B)** UMAP plots showing 20 clusters generated from initial UMAP dimensional reduction and the location of cells within each cluster from each dataset grouping. **C)** Heatmap from unsupervised hierarchical clustering using Pearson's correlation coefficients derived from scaled average expression values of cells within each of the indicated clusters. Cluster number corresponds to UMAP plots above, numbered from top-bottom 0-19.

**Supplementary Figure 6**

Upregulated in SIX1/2miRNA vs. Blastemal (log2 FC > 1.5, adj p <0.05)

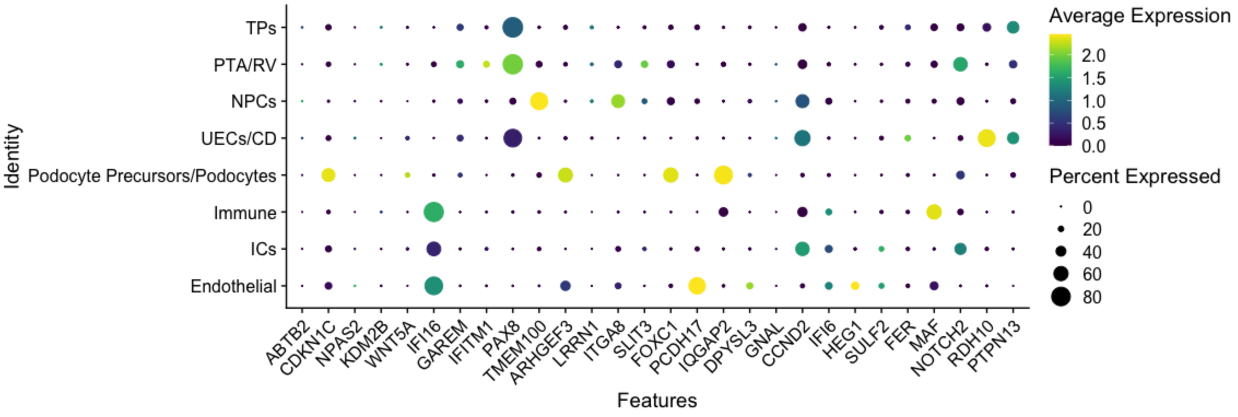


Downregulated in SIX1/2miRNA vs. Blastemal (log2 FC > |1.5|, adj p <0.05)

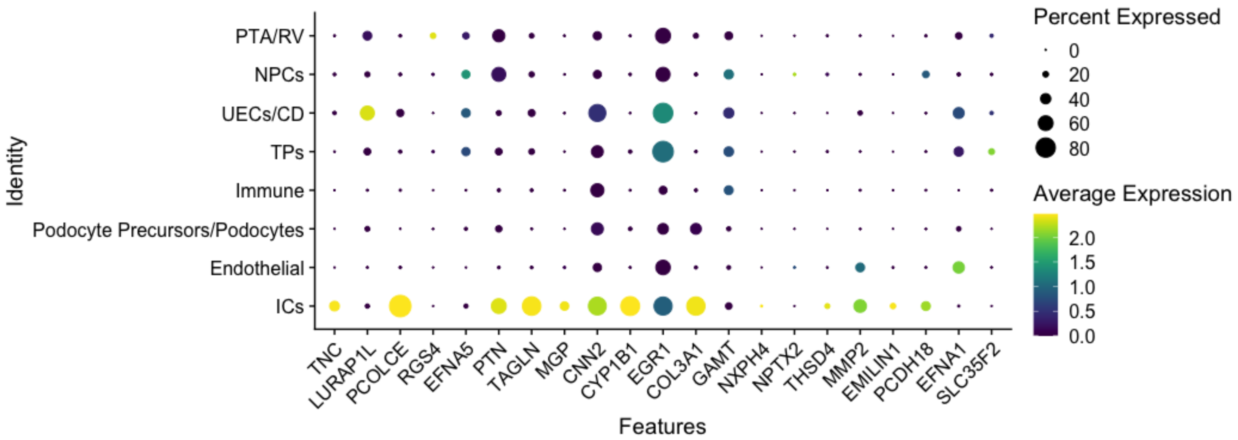


Upregulated in SIX1/2miRNA vs. MIXED/ES (log2 FC > 1.5, adj p <0.05)

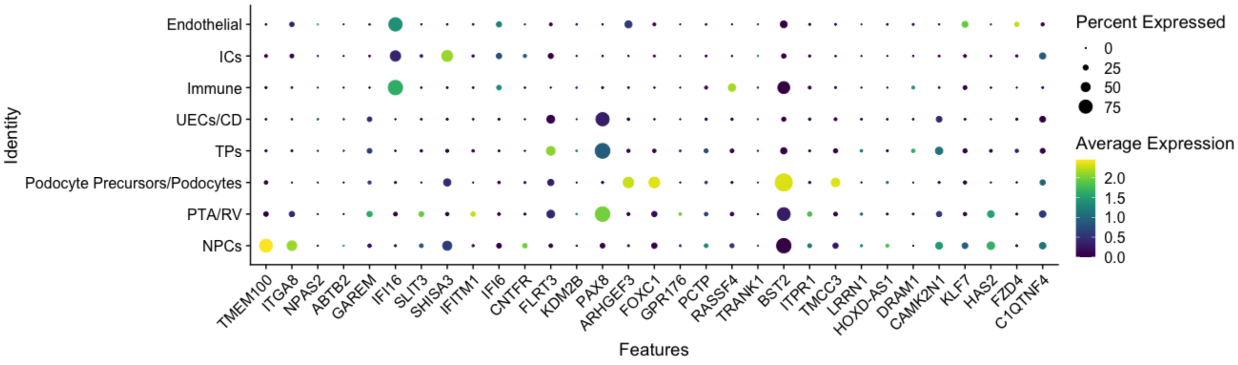

Downregulated in SIX1/2miRNA vs. MIXED/ES (log2 FC > |1.5|, adj p < 0.05)

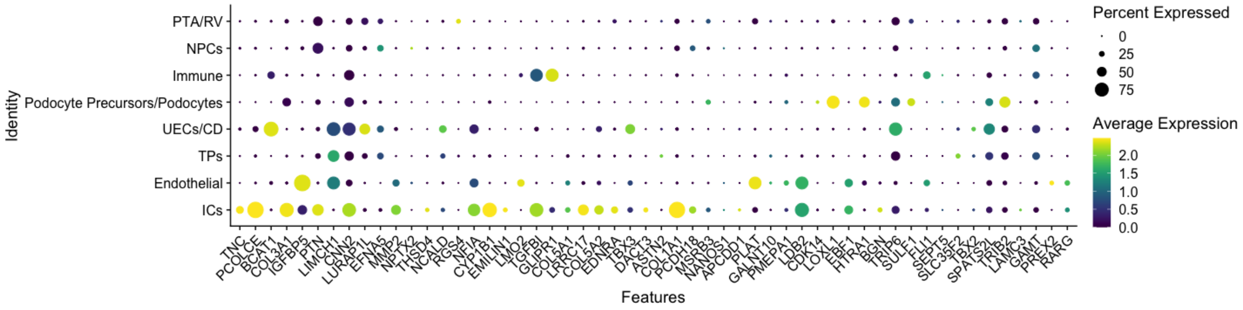


Upregulated and Downregulated in Blastemal vs. MIXED/ES (log2 FC > |1|, adj p < 0.05)

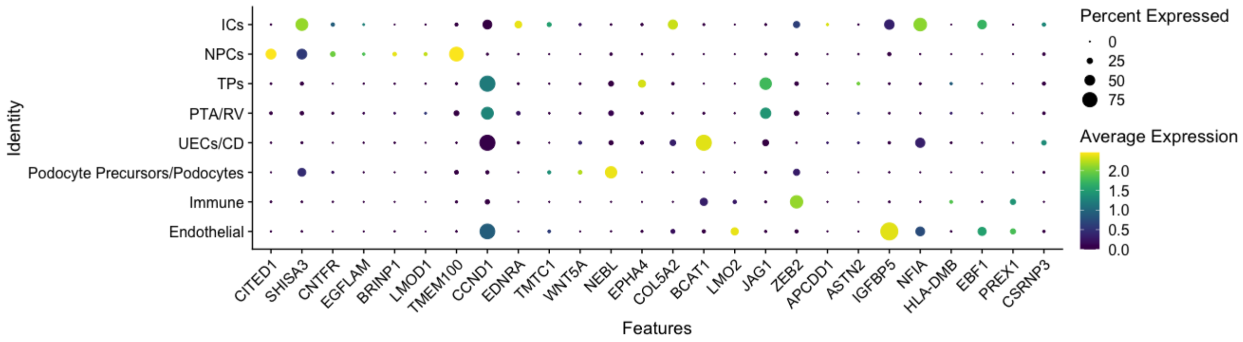


**Supplemental Figure 6:** Dot plots of indicated gene sets displaying the scaled average expression z-score value for each gene on the x-axis and the percent of cells expressing that gene within the cell clusters shown on the y-axis.

**Supplemental Table 1: TARGET Wilms tumor sample IDs and corresponding group assignments used for differential gene expression analysis**

| ***Sample ID*** | ***Group*** | ***Sample ID*** | ***Group*** |
|---|---|---|---|
| **PADZUB** | Blastemal | **PAJLTH** | MIXED/ES |
| **PAEAFB** | Blastemal | **PAJLTI** | MIXED/ES |
| **PAEBXA** | Blastemal | **PAJMEL** | MIXED/ES |
| **PAJLKR** | Blastemal | **PAJMEP** | MIXED/ES |
| **PAJLUJ** | Blastemal | **PAJMKJ** | MIXED/ES |

| | | | |
|---|---|---|---|
| **PAJLWT** | Blastemal | **PAJMKN** | MIXED/ES |
| **PAJMEN** | Blastemal | **PAJMUF** | MIXED/ES |
| **PAJMJT** | Blastemal | **PAJNBN** | MIXED/ES |
| **PAJMMY** | Blastemal | **PAJNCC** | MIXED/ES |
| **PAJMXF** | Blastemal | **PAJNCZ** | MIXED/ES |
| **PAJNCJ** | Blastemal | **PAJNJJ** | MIXED/ES |
| **PAJNGH** | Blastemal | **PAJNLT** | MIXED/ES |
| **PAJNRL** | Blastemal | **PAJNNC** | MIXED/ES |
| **PAJPGY** | Blastemal | **PAJNSL** | MIXED/ES |
| **PAKMSV** | Blastemal | **PAJNTJ** | MIXED/ES |
| **PAKULH** | Blastemal | **PAJNUS** | MIXED/ES |
| **PAKXWB** | Blastemal | **PAJNYT** | MIXED/ES |
| **PAKZFK** | Blastemal | **PAJNZK** | MIXED/ES |
| **PAKZHF** | Blastemal | **PAJNZU** | MIXED/ES |
| **PALDWP** | Blastemal | **PAJPDC** | MIXED/ES |
| **PALGVY** | Blastemal | **PAJPEW** | MIXED/ES |
| **PALKCW** | Blastemal | **PAJPHA** | MIXED/ES |
| **CAAAAB** | DAWT | **PAKECR** | MIXED/ES |
| **CAAAAP** | DAWT | **PAKFME** | MIXED/ES |
| **PAJLIP** | DAWT | **PAKGMU** | MIXED/ES |
| **PAJLKC** | DAWT | **PAKGZX** | MIXED/ES |
| **PAJLPX** | DAWT | **PAKJGM** | MIXED/ES |
| **PAJMKI** | DAWT | **PAKKNS** | MIXED/ES |
| **PAJMLI** | DAWT | **PAKKSE** | MIXED/ES |
| **PAJMLZ** | DAWT | **PAKMUB** | MIXED/ES |
| **PAJMRL** | DAWT | **PAKNAL** | MIXED/ES |

| | | | |
|---|---|---|---|
| **PAJMVU** | DAWT | **PAKNTW** | MIXED/ES |
| **PAJNAA** | DAWT | **PAKRZW** | MIXED/ES |
| **PAJNNR** | DAWT | **PAKSCC** | MIXED/ES |
| **PAJNRH** | DAWT | **PAKSDG** | MIXED/ES |
| **PAJNUP** | DAWT | **PAKVET** | MIXED/ES |
| **PAJNVE** | DAWT | **PAKWPM** | MIXED/ES |
| **PAJNVX** | DAWT | **PAKYFC** | MIXED/ES |
| **PAJNZI** | DAWT | **PAKYLT** | MIXED/ES |
| **PAJNZS** | DAWT | **PAKZER** | MIXED/ES |
| **PAJPAR** | DAWT | **PALERC** | MIXED/ES |
| **PAJPCM** | DAWT | **PALEZT** | MIXED/ES |
| **PAKFYV** | DAWT | **PALFME** | MIXED/ES |
| **PAKNRX** | DAWT | **PALFRD** | MIXED/ES |
| **PAKNXS** | DAWT | **PALGAZ** | MIXED/ES |
| **PAKPDF** | DAWT | **PALGLU** | MIXED/ES |
| **PAKRCC** | DAWT | **PALJIP** | MIXED/ES |
| **PAKUIT** | DAWT | **PALLFB** | MIXED/ES |
| **PAKXXF** | DAWT | **CAAAL** | SIX1/2miRNA |
| **CAAAAC** | MIXED/ES | **CAAAO** | SIX1/2miRNA |
| **CAAAAJ** | MIXED/ES | **PAJMFU** | SIX1/2miRNA |
| **CAAAAS** | MIXED/ES | **PAJMFY** | SIX1/2miRNA |
| **CAAAQ** | MIXED/ES | **PAJMIZ** | SIX1/2miRNA |
| **CAAAR** | MIXED/ES | **PAJMSE** | SIX1/2miRNA |
| **PAECJB** | MIXED/ES | **PAJPAU** | SIX1/2miRNA |
| **PAJLNJ** | MIXED/ES | **PAKRVH** | SIX1/2miRNA |
| **PAJLSP** | MIXED/ES | **PALDTE** | SIX1/2miRNA |

#### Supplemental Table 2: Oligo and primer sequences used in cloning, luciferase assays, EMSAs, and PBMs

| |
|---|
| **>SIX1_enhancer_Luciferase assay** |
| ATTAGCTAGCCCAGCCGCGGCCCCAGCCCTCCCCCCAGCCTGTGCTGGGCTCCGCTTTCCCTCCATCAACTCCAAGCCGAATTCAATCCGAGAAGGCTCC<br>TTTGAGCTTTTGTGTTTGCTGGGGGAGATGTGGGCGCAGGAGGGATCGCGTTACAACTTTCATTTCCTGAAATGTTTGAGGGAACATCCAGGGTTTTTATC<br>CCCACATCAGGCCGGGCGATGGGCTCGAGTTTCAGGCCTTGTCACTCAGCTGTCACCAAACAAACGAAGCTCTCAGAGCCCAGGAGAGGGAGAGCTACC<br>TGCTATTCATGACCCCTGGAGCAGGTGATCGCTCATGGGAAAAACAGGTAGAATTAATCATAGGACTGTTCTCTGTTTCCTCTCCTTTTTGGCAGACCTGCC<br>CACAGTGCGAAACCTATCAGCAAACAAATTAACTCTTTCTTGTCACCCAGGGGAATTAAAAATACGGTTTTAGAAAATAACTAAAACACACACGTTTCTCACC<br>AAATGACAAAATGGGAGTTGGAAGGAACATCTTACATCCCGCCCTACCTTGCAATTTCTTAGTTTGTGGGATTCTCATCTTGGTAGGATTTAGAACTTGGGG<br>AAACTGGTAGAGAATGAGGCATTCTCATGTTACCTGGTCCTACTACTGGATGGAGCCCCAGGTGTGGTCCCTGAACTCAGGTAGATTTCAACATCTCCTTCT<br>TGGCTGGGTGTCAATGCAGTTGGGCTTGATCCTACTAAAACAGAAAATCTCTACTCCAACAGGAATTTCTAAGAAATTTCTGTAAATTAGTGTTTTGCTTCCC<br>CCAGCCTCTTCCCTAAGTCAATCATGTTGAGCAAGCTTAATA |
| **>WNT5A_promoter_Luciferase assay** |
| TTATACCCAGCTAGCACGTAAGTCTAACCCTGCCGCACTGCATCGCCCATAGCCCCTGAAGGAGCCCCTTCCACAGAAAAGAAAAGAAAGGTGAGCCTCTT<br>TAAGGCGGTGGAAGAGCCTGGCTTGGAACCTGTGCCGTAAAAGGGGCAGAGGGGACCTAGGCAGCCCTGGTAAAGCTATGGGGCTCAGGGGCGTGCCA<br>AGGTTTTTCTCCGTGAGCCGCCCCTTTGGCCTGGACGCTTCGGGGCTTCTCAAAGAGGAAATGCTTATGTGGTCCCCAGCGCCTGCTAAGCAGGGCTCCA<br>CACCCAAGGCCCAGTTGTCCCCAAAACGCTGCAAAGCTGGGGGGCGCATCCTGGAGAATGGAAATCTGGGGTTTCCCCAGCTAGGAGAGAGAAGGCTCC<br>GGCTATCTCCCCACCCCCGCCCTAAGTGTCAAATTTCCTCCAGGGGAGGGAGTGGGCTGCAAAGTCTGCTTCTCGCGCAGCCCAGGCTGCAAAGTCAACT<br>CTCCCCAAGGGCAGCCGATGCCGCGTGCACACACATCATACACATTCACACTCGTGCACATTTACACACTCACACGCGTGCATAGACACACACGCTGCGA<br>CATGTTTCCGAGTCAGCGGCCCAGATTGGTGCTGGCCGCGTGCACTTTCCAAGCTTCGCGGCGAGCGGGGCGCGTGGGGCGGGGCTCAAGCAGCAGAG<br>AAATTGATAACAGATTCGGCGGATTACAGCGGATCTCTTTGTTAGAGCCGAAGCCACACAAACCGAACCCACTCCCAGCCCGAAGCCCCCAGGGAGAGTC<br>CACCAGTTCCCAGAGCCCAAAGCTTAATAGC |
| **>WNT5A_distal1_Luciferase assay** |
| TTATACCCAGCTAGCGTTTTATTAATTCCATGCTGGAGTTCCATAGCTGGGCCAGTCGGTTTGTAAGTTACCTCTGGAACAGTAGCTGAACTACCCAGAGTG<br>CATGATTGGAAAAGAGGAAATGGGGGAGGCTAGTCCTCCTCAACCTCTCACAGGAGAGAAATATGGTTTTCTCCAGCTCCTACAGACAGGTTCCTGGGGCT<br>ATCTGGGGCTGGAGCCTCTAGGCAAGCTGATTTGAAACGCATCCAAGCGGATATCTTGCTTCAAAGAGTCAGGCCCCCCCGGGGCAGAGCCTGAACCTGA<br>GTACTGGAAGAGGACCAGGGGGCTGGAATTCAGGCACTGCCCTCCCATCCCCCAGGCCCGGGAGGGTTTGTTTAACCCCTCAGGGGTGCTGTGTGCCTT<br>CCTTGTCGAGGTGGTTTGCGGCTTCGTGAAGCGATATTTATAGAGTGCTCTGTGATTCTTGGGTGAACATTCGTTTTATAAATGAATGATCAAATTTATTAGC<br>AAAGAGTAACTAACCCGTCTACATTTATTTTGTTGATAATACAAATTTACAAGGTTTTAATTGCCAGTTTTAAAACCCCCAAAACCCATAAATAAATACACGGG<br>TAATAAAATATCTTCTCACTGTGCCCTTGACTTCTGTAGTTACATGAGGGCTTATGGGAATGTCTGAAGGAGCTTTCAGATGGAATCACAAAACTTGTTTGAA<br>AGGACTCAGGGCGGGAAACAGTCTAATTCAAACCTGGCTGGTGGTAATTTCATGTATGTTAACCAAAATGCTACTACTGATCATGAAGCTTAATAGC |
| **>MEIS1-3XFLAG_Fwd primer for cloning** |
| GATCAGGATCCGCCGCCACCATGGCGCAAAGGTACGACGATCTA |
| **>MEIS1-3xFLAG_Rev primer for cloning** |
| GATCACTCGAGTTACTTGTCATCGTGATCCTTGTAATCGATGTCATGATCTTTATAATCACCGTCATGGTCTTTGTAGTCCATGTAGTGCCACTGCCCCTCCA<br>TGC |
| **>WNT5A_distal2_Luciferase assay** |
| TTATACCCAGCTAGCGTGTACACACACGGTCGTGTTACAATTCTCATTTTGCAAAGTTTATGCAAAAACCAAACACCTGGGTTCAGAGTTTCCTAAAGGAGT<br>CATCTGAAGTAGGTGCTTTACGCCAAAACGTCACAAAGATTTATGTGCTTTCATTTGTGCATTAAATTGAGGACAGGTGGGAGAATGCTCAGGCCTGAGAAA<br>AACTGATAGCTCATTTCTCCCTTCGAAGAGAGATGGCTGTTATGACTACTGCTGGTTTAGATAAAATAGATACAGACTTTGTTTAAAAAAAAAAAAGAGGGCT<br>CATGTTTGAAAAACAGTATTTCAGCAGTCAAATAATTACCTCTGCGATCATGTTTCTCACAAATGGAAAACCCTGGACTGAGATCCCACAGTGAGTCCTGCT<br>CCTAGTTCAAACAACAGGCAATACCATGCCCAACAGCCAAGAAAATGGCCGGACCTCCCTTCACACTTGCTGAGGAAGGGTCCCTGGAATTGAGGCAAAT<br>GGTGGCCTCAACACATTCCATTTTAATGGCTTGCAACAGAGTCAGTTACACATATGTAACCACTCACTTTTTAATTTGATTCTTGTTTCAAACACCTTTTCAGG<br>ACCCAATACATCTAAAAAATGTCATCACTTGATAGGTCGCTACTCTATCCCTGGCCTTGAAACAAGTGATTCGATGTGCTGAGTATTCACCAGGATTTGAAAT<br>AAGACTGTGAATTTGCATTTCAGAAAGGGAATACTATTTGGAAAGCCCAGTCTGCCAGACTTTTCACAAAGGGAAAGGAGAGCTCCACTGGGAAAGCCTGC<br>TGGTCAGCCAAGCTTAATAGC |
| **>WNT5A_promoter_EMSA** |
| CCCGCCCTAAGTGTCAAATTTCCTCCAGGGGAG |
| GGGCGGGATTCACAGTTTAAAGGAGGTCCCCTC |
| **>WNT5A_promoter_EMSAmut** |
| CCCGCCCTAAGT**A**TCAAATTTCCTCCAGGGGAG |
| GGGCGGGATTCA**T**AGTTTAAAGGAGGTCCCCTC |
| **>SIX1_gBlock for protein purification and luciferase assay** |
| GTTGTTCTCGAGGCCGCCACCATGTCGATGCTGCCGTCGTTTGGCTTTACGCAGGAGCAAGTGGCGTGCGTGTGCGAGGTTCTGCAGCAAGGCGGAAAC<br>CTGGAGCGCCTGGGCAGGTTCCTGTGGTCACTGCCCGCCTGCGACCACCTGCACAAGAACGAGAGCGTACTCAAGGCCAAGGCGGTGGTCGCCTTCCAC<br>CGCGGCAACTTCCGTGAGCTCTACAAGATCCTGGAGAGCCACCAGTTCTCGCCTCACAACCACCCCAAACTGCAGCAACTGTGGCTGAAGGCGCATTACG<br>TGGAGGCCGAGAAGCTGCGCGGCCGACCCCTGGGCGCCGTGGGCAAATATCGGGTGCGCCGAAAATTTCCACTGCCGCGCACCATCTGGGACGGCGAG<br>GAGACCAGCTACTGCTTCAAGGAGAAGTCGAGGGGTGTCCTGCGGGAGTGGTACGCGCACAATCCCTACCCATCGCCGCGTGAGAAGCGGGAGCTGGC<br>CGAGGCCACCGGCCTCACCACCACCCAGGTCAGCAACTGGTTTAAGAACCGGAGGCAAAGAGACCGGGCCGCGGAGGCCAAGGAAAGGGAGAACACCG<br>AAAACAATAACTCCTCCTCCAACAAGCAGAACCAACTCTCTCCTCTGGAAGGGGGCAAGCCGCTCATGTCCAGCTCAGAAGAGGAATTCTCACCTCCCCAA<br>AGTCCAGACCAGAACTCGGTCCTTCTGCTGCAGGGCAATATGGGCCACGCCAGGAGCTCAAACTATTCTCTCCCGGGCTTAACAGCCTCGCAGCCCAGTC<br>ACGGCCTGCAGACCCACCAGCATCAGCTCCAAGACTCTCTGCTCGGCCCCCTCACCTCCAGTCTGGTGGACTTGGGGTCCTAAGGATCCTTGTTG |
| **>SIX1-Q177R_gBlock for protein purification and luciferase assay** |
| GTTGTTCTCGAGGCCGCCACCATGTCGATGCTGCCGTCGTTTGGCTTTACGCAGGAGCAAGTGGCGTGCGTGTGCGAGGTTCTGCAGCAAGGCGGAAAC<br>CTGGAGCGCCTGGGCAGGTTCCTGTGGTCACTGCCCGCCTGCGACCACCTGCACAAGAACGAGAGCGTACTCAAGGCCAAGGCGGTGGTCGCCTTCCAC<br>CGCGGCAACTTCCGTGAGCTCTACAAGATCCTGGAGAGCCACCAGTTCTCGCCTCACAACCACCCCAAACTGCAGCAACTGTGGCTGAAGGCGCATTACG<br>TGGAGGCCGAGAAGCTGCGCGGCCGACCCCTGGGCGCCGTGGGCAAATATCGGGTGCGCCGAAAATTTCCACTGCCGCGCACCATCTGGGACGGCGAG<br>GAGACCAGCTACTGCTTCAAGGAGAAGTCGAGGGGTGTCCTGCGGGAGTGGTACGCGCACAATCCCTACCCATCGCCGCGTGAGAAGCGGGAGCTGGC<br>CGAGGCCACCGGCCTCACCACCACCCAGGTCAGCAACTGGTTTAAGAACCGGAGGAGAAGAGACCGGGCCGCGGAGGCCAAGGAAAGGGAGAACACCG<br>AAAACAATAACTCCTCCTCCAACAAGCAGAACCAACTCTCTCCTCTGGAAGGGGGCAAGCCGCTCATGTCCAGCTCAGAAGAGGAATTCTCACCTCCCCAA<br>AGTCCAGACCAGAACTCGGTCCTTCTGCTGCAGGGCAATATGGGCCACGCCAGGAGCTCAAACTATTCTCTCCCGGGCTTAACAGCCTCGCAGCCCAGTC<br>ACGGCCTGCAGACCCACCAGCATCAGCTCCAAGACTCTCTGCTCGGCCCCCTCACCTCCAGTCTGGTGGACTTGGGGTCCTAAGGATCCTTGTTG |
| **>EYA1-2xHA_Fwd primer for cloning** |
| GTTGTTGAATTCGCCGCCACCATGGAAATGCAGGATCTAACCA |
| **>EYA1-2xHA_Rev primer for cloning** |
| GTTGTTCTAGATTAAGCGTAATCTGGAACATCGTATGGGTAAGCGTAATCTGGAACATCGTATGGGTACAGGTACTCTAATTCCAAGG |
| **CONTINUED ON NEXT PAGE** |

| **>SIX1_nGST_HD for PBM** |
|---|
| CATATGATGTCCCCTATACTAGGTTATTGGAAAATTAAGGGCCTTGTGCAACCCACTCGACTTCTTTTGGAATATCTTGAAGAAAAATATGAAGAGCATTTGT<br>ATGAGCGCGATGAAGGTGATAAATGGCGAAACAAAAAGTTTGAATTGGGTTTGGAGTTTCCCAATCTTCCTTATTATATTGATGGTGATGTTAAATTAACACA<br>GTCTATGGCCATCATACGTTATATAGCTGACAAGCACAACATGTTGGGTGGTTGTCCAAAAGAGCGTGCAGAGATTTCAATGCTTGAAGGAGCGGTTTTGG<br>ATATTAGATACGGTGTTTCGAGAATTGCATATAGTAAAGACTTTGAAACTCTCAAAGTTGATTTTCTTAGCAAGCTACCTGAAATGCTGAAAATGTTCGAAGAT<br>CGTTTATGTCATAAAACATATTTAAATGGTGATCATGTAACCCATCCTGACTTCATGTTGTATGACGCTCTTGATGTTGTTTTATACATGGACCCAATGTGCCT<br>GGATGCGTTCCCAAAATTAGTTTGTTTTAAAAAACGTATTGAAGCTATCCCACAAATTGATAAGTACTTGAAATCCAGCAAGTATATAGCATGGCCTTTGCAG<br>GGCTGGCAAGCCACGTTTGGTGGTGGCGACCATCCTCCAAAATATCGGGTGCGCCGAAAATTTCCACTGCCGCGCACCATCTGGGACGGCGAGGAGACC<br>AGCTACTGCTTCAAGGAGAAGTCGAGGGGTGTCCTGCGGGAGTGGTACGCGCACAATCCCTACCCATCGCCGCGTGAGAAGCGGGAGCTGGCCGAGGC<br>CACCGGCCTCACCACCACCCAGGTCAGCAACTGGTTTAAGAACCGGAGGCAAAGAGACCGGGCCGCGGAGGCCAAGGAAAGGGAGAACACCGAAAACA<br>ATAACTCCTCCTCCAACTAACTCGAG |
| **>SIX1-Q177R_nGST_HD for PBM** |
| CATATGATGTCCCCTATACTAGGTTATTGGAAAATTAAGGGCCTTGTGCAACCCACTCGACTTCTTTTGGAATATCTTGAAGAAAAATATGAAGAGCATTTGT<br>ATGAGCGCGATGAAGGTGATAAATGGCGAAACAAAAAGTTTGAATTGGGTTTGGAGTTTCCCAATCTTCCTTATTATATTGATGGTGATGTTAAATTAACACA<br>GTCTATGGCCATCATACGTTATATAGCTGACAAGCACAACATGTTGGGTGGTTGTCCAAAAGAGCGTGCAGAGATTTCAATGCTTGAAGGAGCGGTTTTGG<br>ATATTAGATACGGTGTTTCGAGAATTGCATATAGTAAAGACTTTGAAACTCTCAAAGTTGATTTTCTTAGCAAGCTACCTGAAATGCTGAAAATGTTCGAAGAT<br>CGTTTATGTCATAAAACATATTTAAATGGTGATCATGTAACCCATCCTGACTTCATGTTGTATGACGCTCTTGATGTTGTTTTATACATGGACCCAATGTGCCT<br>GGATGCGTTCCCAAAATTAGTTTGTTTTAAAAAACGTATTGAAGCTATCCCACAAATTGATAAGTACTTGAAATCCAGCAAGTATATAGCATGGCCTTTGCAG<br>GGCTGGCAAGCCACGTTTGGTGGTGGCGACCATCCTCCAAAATATCGGGTGCGCCGAAAATTTCCACTGCCGCGCACCATCTGGGACGGCGAGGAGACC<br>AGCTACTGCTTCAAGGAGAAGTCGAGGGGTGTCCTGCGGGAGTGGTACGCGCACAATCCCTACCCATCGCCGCGTGAGAAGCGGGAGCTGGCCGAGGC<br>CACCGGCCTCACCACCACCCAGGTCAGCAACTGGTTTAAGAACCGGAGGAGAAGAGACCGGGCCGCGGAGGCCAAGGAAAGGGAGAACACCGAAAACA<br>ATAACTCCTCCTCCAACTAACTCGAG |
| **>Minimal_Promoter1 for pBV-Luc** |
| AGTAAGCTTGGGGGTATATAATGGATCCGGTATCGAGATCTGCGATCTAAGTAAGTTGGCATTCCGGTACTGTTAAAGCCACCATGGAAC |
| **>Minimal_Promoter2 for pBV-Luc** |
| GTTCCATGGTGGCTTTAACAGTACCGGAATGCCAACTTACTTAGATCGCAGATCTCGATACCGGATCCATTATATACCCCCAAGCTTACT |
